# Hierarchical global and local auxin signals coordinate cellular interdigitation in *Arabidopsis*

**DOI:** 10.1101/2024.06.17.599171

**Authors:** Patricio Pérez-Henríquez, Hongjiang Li, Xiang Zhou, Xue Pan, Wenwei Lin, Wenxin Tang, Shingo Nagawa, Deshu Lin, Tongda Xu, Marta Michniewicz, Michael J. Prigge, Lucia C. Strader, Mark Estelle, Ken-ichiro Hayashi, Jiří Friml, Linlin Qi, Zhongchi Liu, Jaimie Van Norman, Zhenbiao Yang

## Abstract

The development of multicellular tissues requires both local and global coordination of cell polarization, however, the mechanisms underlying their interplay are poorly understood. In Arabidopsis, leaf epidermal pavement cells (PC) develop a puzzle-piece shape locally coordinated through apoplastic auxin signaling. Here we show auxin also globally coordinates interdigitation by activating the TIR1/AFB-dependent nuclear signaling pathway. This pathway promotes a transient maximum of auxin at the cotyledon tip, which then moves across the leaf activating local PC polarization, as demonstrated by locally uncaged auxin globally rescuing defects in *tir1;afb1;afb2;afb4;afb5* mutant but not in *tmk1;tmk2;tmk3;tmk4* mutants. Our findings show that hierarchically integrated global and local auxin signaling systems, which respectively depend on TIR1/AFB-dependent gene transcription in the nucleus and TMK-mediated rapid activation of ROP GTPases at the cell surface, control PC interdigitation patterns in Arabidopsis cotyledons, revealing a mechanism for coordinating a local cellular process with the development of whole tissues.

## Introduction

Cell polarization along the plane of an organ’s surface, known as planar cell polarity (PCP), must coordinate signaling at two different functional levels; locally between adjacent cells and globally across the entire tissue ^1^. Despite the critical importance of PCP in various developmental processes in animals and human health ^2,3^, it remains poorly understood how the two scales of signaling are coordinated and linked to the regulation of tissue and organ development. In animals, including humans, PCP is modulated by peptidyl WNT signals via both cytoplasmic and nuclear signaling pathways ^4^. In plants, a key signal controlling pattern formation and morphogenesis is auxin ^5^. Auxin is polarly transported across cells, primarily via the PIN family of auxin transporters ^6^, giving rise to concentration gradients essential for many developmental processes such as establishment of the planar polarity during root hair initiation ^7,8^. A long-range auxin gradient along the root was found to coordinate with a short-range auxin signal that promotes root hair initiation, but the underlying signaling pathways are not known ^7,8^. Auxin signal transduction is mediated by two main perception modules. The nuclear TRANSPORT INHIBITOR RESPONSE 1/ AUXIN-SIGNALING F-BOX (TIR1/AFB) module regulates nuclear gene expression ^9^, whereas a non-canonical perception module, which relies on the auxin-binding protein 1 (ABP1) and ABP1-like (ABL) proteins and their interacting partners TRANSMEMBRANE KINASES (TMKs), typically regulates plasma membrane activities and cytoplasmic responses ^10–16^. Whether and how these two auxin signaling modules coordinately regulate a given auxin-mediated process remains unknown ^17^. Here, we investigate their functional relationship and connections to local and global coordination of cell polarization during the planar interdigitation to form the puzzle-piece-shaped pavement cells (PCs) in *Arabidopsis* embryonic leaves (cotyledons) ^18,19^. Our previous studies suggest that TMK-perceived auxin locally coordinates PC interdigitation by regulating Rho GTPase-based signaling pathways, leading to cytoskeletal re-organization and planar cell polarization ^10,12,20–23^. In this study, we show that nuclear TIR1/AFB auxin receptors globally coordinate PC interdigitation throughout the cotyledon via the transcription-based auxin signaling pathway and act upstream of the TMK module by generating an auxin signal that activates TMKs. Thus, we propose a hierarchical self-organizing signaling system that controls pattern formation in *Arabidopsis* cotyledons by integrating the local, cellular-level coordination of cell polarity with its global coordination at the tissue level. This design principle may be analogous to the regulation of pattern formation by WNT signaling in animals, which involves gene activation as well as Rho GTPase-dependent signaling.

## Results

### The spatiotemporal wave of PC interdigitation correlates with the dynamic auxin distribution with a maximum at the cotyledon tip

To understand the global coordination of PC interdigitation, we monitored PC shape in the adaxial epidermis of expanding embryonic leaves (cotyledons) at 0, 24, 36, and 72 <u>h</u>ours <u>a</u>fter <u>p</u>lating seeds (HAP). Software-assisted quantification ^24^ of the Margin Roughness (MR), which accounts for early emerging lobes otherwise undetectable by currently automated approaches ^24–26^ (**Figure 1A**), revealed a dominant presence of cells with little or no lobes at 0 and 24 HAP (prior to seed germination) (**Figure 1B, C**). Then, PC interdigitation was initiated coinciding with germination, spreading from the apical region to upper-mid regions at 36 HAP (**Figure 1B, C, S1A**) and finally reaching the base of the cotyledon at 72 HAP (**Figure 1B, C**). These PC shape changes imply the existence of some global developmental signal(s), starting at the tip and spreading to the remaining parts of the cotyledon.

**Figure 1.**
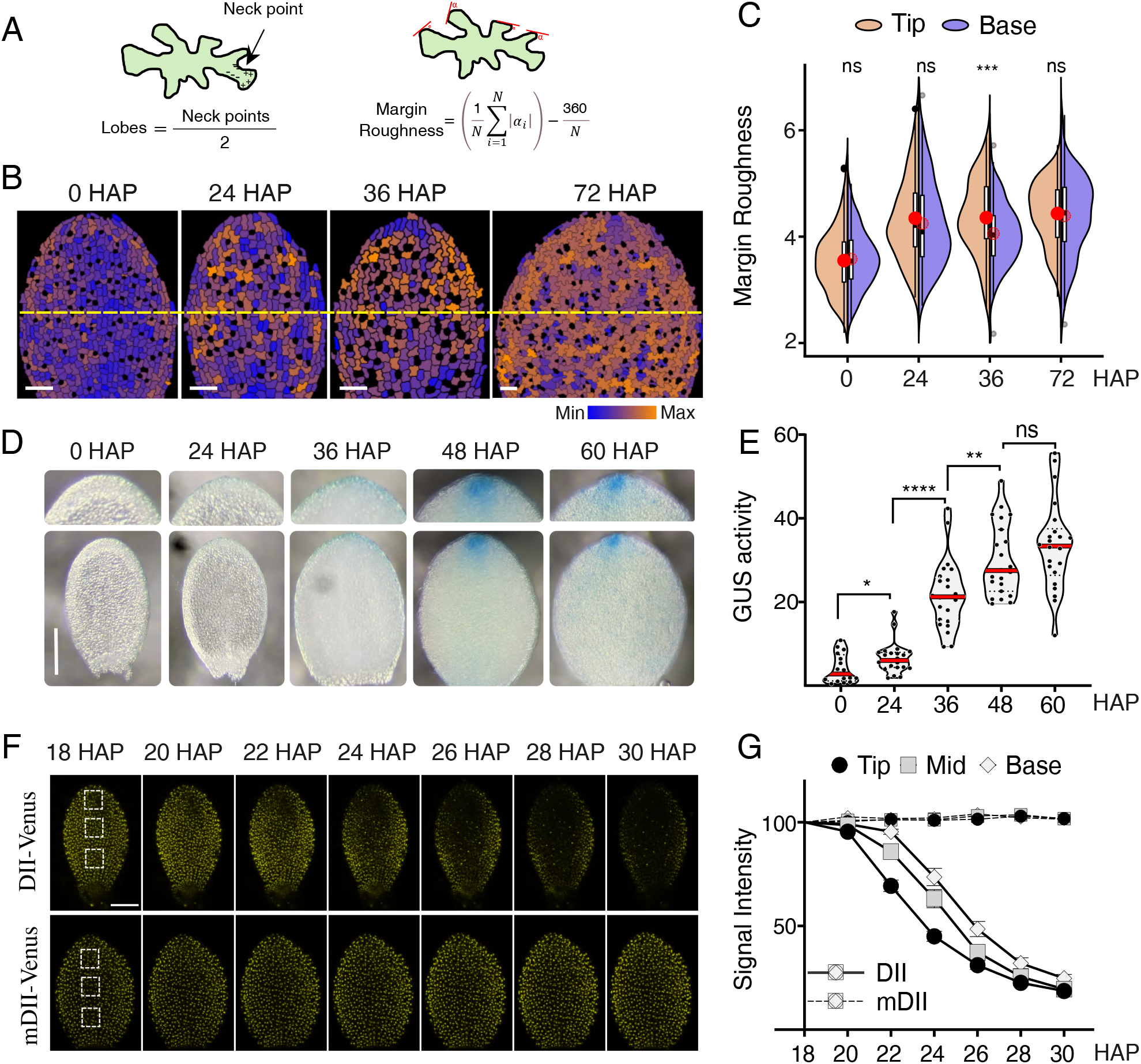
The progressive activation of pavement cell (PC) interdigitation follows a similar pattern of increase in auxin levels that begins at the tip of young Arabidopsis cotyledons. (A) Schematic of PC metrics quantification for margin roughness (MR) and lobe count (Lobes). (B) Heatmap of MR shows that PC interdigitation first occurs in the tip and progressively spreads to the middle and basal regions of expanding cotyledons. At the indicated hours after plating (HAP), wild-type (Col-0) cotyledon PCs were imaged using laser scanning confocal microscopy, and the degree of MR was computed per cell and color-coded as shown in the color scale. Scale bars = 50 μm. Yellow dashed lines separate the top and bottom half of the early expanding cotyledons. (C) Quantification of MR of pavement cells at the cotyledon’s base and tip, defined as the top and bottom half of early expanding cotyledons, analyzed with the software PaCeQuant ^24^. Cell borders were obtained by staining with propidium iodide. Cotyledons were dissected before imaging: swollen seed (0 HAP), ruptured seed testa (24 HAP), emerged radicle (36 HAP), greening cotyledons (48 HAP), green opening cotyledons (60 HAP) and green open flat globular cotyledons (72 HAP). Box plot inside each violin plot depicts four quartiles and the median. Red dot depicts the average. n=231-368 cells, *t*-test ***p<0.001. (D) GUS histochemical assay in the cotyledons of a *DR5::GUS* line suggests an apparent tip-high auxin maximum at 24 HAP, a clear apical margin-high maximum at 36 HAP, and a conspicuous tip-high maximum at 48 and 60 HAP. Scale bar = 150 μm. (E) This is confirmed by GUS activity quantification in cotyledons at the same developmental time points shown in D by fluorometric detection of 4-methylumbelliforone (4-MU), n=24 cotyledons, *t*-test, *p<0.05, **p<0.01, ****p<0.0001. Note that GUS activity at 24 HAP was significantly higher than at 0 HAP. (F) Representative images from a time-lapse of cotyledons in plants expressing DII-Venus (upper row) or mDII-Venus (lower row, a mutation in DII that makes it insensitive to auxin). (G) Quantitative analysis of Venus signal intensity in cells on the tip, middle, and base of cotyledons, defined as shown by the dashed boxes in F, 18 HAP. In DII-Venus (DII) cotyledons, tip cells (black) show a reduction in signal intensity as early as 22 HAP. Reduction of signal intensity was then observed in cells in the middle (gray) and, finally, in the base (white). In contrast, signal intensity was unchanged in mDII-Venus (mDII) cotyledon for all regions. Plot shows mean + standard error. n=20-22 cotyledons, each from different seedlings from 3 experimental replicates.

We speculated that auxin might be such a global coordinator of PC interdigitation as it is the major morphogenetic signal, which forms concentration gradients and/or maxima ^27^ and is required and sufficient to promote PC interdigitation ^20^. Thus, we examined *DR5::GUS* expression, which reports auxin-responsive gene transcription. At 24 HAP (prior to the initiation of PC interdigitation), *DR5::GUS* expression was first detected at the tip and marginal regions close to the tip of cotyledons, and at 36 HAP, the apical GUS signal became stronger. At 48-60 HAP GUS signal spreaded to a larger area from the tip, suggesting an auxin maximum at the tip (**Figure 1D**). The dynamic changes in GUS activity were corroborated by quantification of GUS activity in a fluorometric assay (**Figure 1E**). A tip-high auxin maximum is consistent with direct auxin measurements in tobacco leaves, which show an auxin maximum at the tip of the youngest leaves ^28^. Furthermore, imaging of DII-Venus, an auxin reporter based on the auxin-induced degradation of the DII domain found in the AUX/IAA transcriptional repressors ^29^, suggested the existence of a transient auxin maxima along the proximodistal axis of the cotyledon (**Figure 1F, G**). The reverse DII-Venus gradient was quite evident between 22 and 28 HAP (**Figure 1G**). After that, the DII-Venus signal was weak and no longer exhibited a gradient or maximum (**Figure 1F, G**). Altogether, we concluded that the progressive activation of PC interdigitation from the cotyledon tip towards the base is preceded by the formation of a transient auxin maximum at the tip of cotyledons.

### The auxin maximum and dynamic wave of PC interdigitation is modulated by cytokinin

Both *DR5::GUS* and reverse DII-Venus signals indicate the existence of dynamic tip-high auxin maxima. However, the two reporters clearly exhibited different dynamics. The former persisted beyond 36 HAP, whilst the latter was very transient, occurring between 22 and 28 HAP. The difference might be explained by the nature of these two different reporters: DII-Venus more directly reflects the input of auxin concentrations ^29^, whereas *DR5::GUS* indicates more downstream transcriptional output, thus also integrating auxin-independent signals such as cytokinins or brassinosteroids ^30,31^.

Cytokinin acts in a manner opposite to auxin in many developmental processes ^32,33^ and suppresses PC interdigitation acting upstream of ROP signaling ^34^. Thus, we hypothesized that cytokinin may suppress auxin-induced gene transcription explaining the difference between the *DR5::GUS* and DII-Venus reporters. Consistently, the cytokinin signaling marker *ARR5::GUS* ^35^ was excluded only from the apical and partially excluded from the marginal regions, an expression pattern complementary to *DR5::GUS* (**Figure S1B**). Indeed, the over-activation of cytokinin signaling by *ARR20* overexpression dramatically reduced *DR5::GUS* expression in young cotyledons (**Figure S1B**). In contrast, *DR5::GUS* was ectopically activated throughout the un-germinated or 24-HAP cotyledons, when cytokinin signaling was blocked by *ARR7* overexpression or in the *ahk3 cre1* mutant, which lacks the two redundant cytokinin receptors AHK3 and CRE1 (**Figure S1C**). Ectopic *DR5::GUS* expression was associated with the premature activation of PC interdigitation throughout the cotyledon in un-germinated seeds (**Figure S1C**). Furthermore, exogenous auxin treatment increased PC interdigitation equally in Col-0 as in *ARR20*-OX plants (**Figure S1D, E**). These results suggest cytokinin signaling acts as a developmental brake to prevent premature activation of PC interdigitation in un-germinated cotyledon by suppressing nuclear auxin responses.

Exogenous cytokinin treatment restricts the TIR1/AFB-based nuclear transcriptional auxin responses to the tip and margin of cotyledons at 24-72 HAP (**Figure S1F**). However, at this stage, endogenous cytokinin signaling did not suppress the tip-to-base progressive activation of PC interdigitation although it suppresses *DR5::GUS* expression in the center and base (**Figure 1C, D**). These results not only revealed a role of cytokinin in restricting the transcriptional auxin response gradient mediating PC interdigitation but also suggest that the PC interdigitation process in individual cells may be directly activated by auxin coming from the cotyledon tip.

### Ectopically-generated local auxin maxima induce global changes in PC interdigitation

Based on the above observations, we hypothesized the auxin maximum at the tip of cotyledons acts globally to promote PC interdigitation throughout the entire cotyledon surface. To test this, we first conducted surgical removal of cotyledon tips from 24 HAP seedlings, and quantified interdigitation after 1 day. The mTalin-GFP expression indicated that cotyledons remained viable after tip removal (**Figure S2A**). Tip removal greatly inhibited PC interdigitation in the central parts of cotyledons (**Figure S2A, B**), supporting the importance of the auxin maximum at the cotyledon tip.

Given tight regulation of spatial distribution of auxin in plants, we were interested in whether the auxin maximum at the cotyledon tip acts globally to promote PC interdigitation. For this, we created ectopic local auxin maxima using UV light-sensitive caged auxins [caged NAA and IAA, (2,5-dimethoxyphenyl)(2-nitrobenzyl), DMPNB-NAA and DMPNB-IAA, respectively] **(Figure 2A, S2C**). Carefully calibrated UV radiation generated no apparent cell damage or background fluorescence (**Figure S2D**). Localized UV irradiation generated a local auxin increase measured in 24 HAP cotyledons of the ratiometric auxin reporter R2D2 (**Figure 2B, C).** Nanomolar auxin concentrations were used for these experiments aimed to detect rapid protein degradation. Uncaged auxin is detectable even at a single-cell resolution (**Figure S2E**), and is consistent with previous reports in tobacco cells ^36,37^. Furthermore, the TIR1/AFBs inhibitor auxinole blocked the response to uncaged auxin ^38^, ruling out non-specific effects as the cause of DII-Venus degradation (**Figure S2E**). Importantly, *DR5::GFP* expression was induced in the entire cotyledon within 20 hours after the UV treatment (**Figure S2F)**.

**Figure 2.**
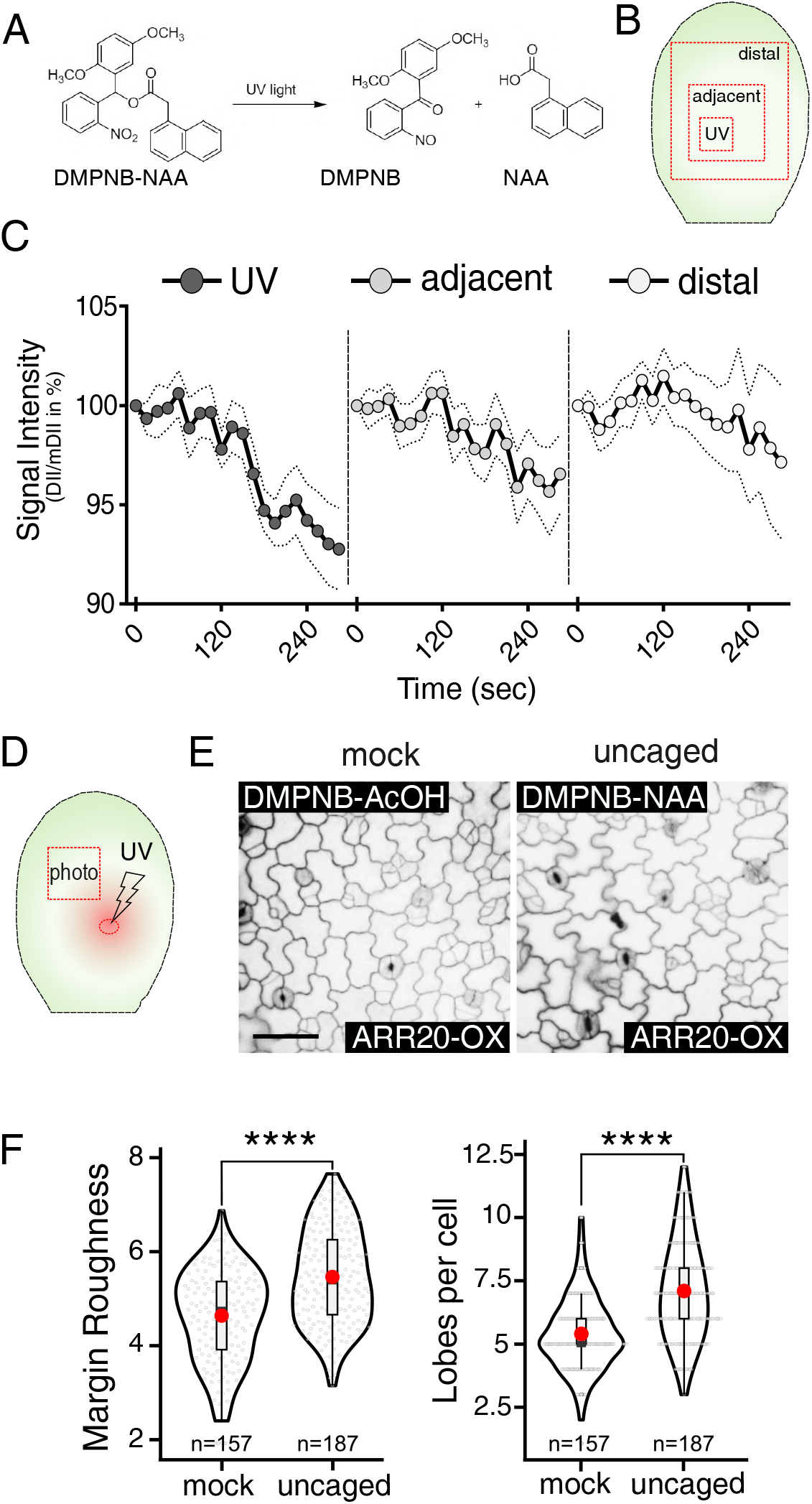
Ectopic local auxin maximum globally activates PC interdigitation. (A) Auxin uncaging reaction. UV light breaks caged DMPNB-NAA/IAA into uncaged active auxin and the cage, see also Figure S2C. UV treatment of DMPNB-AcOH (mock) allows the release of acetic acid to emulate auxin acidity without auxin response. (B) Schematic representation of auxin uncaging experiment testing the efficacy of uncaging in the UV-treated area and the adjacent, and more distal areas. (C) Efficacy of auxin uncaging by quantification of the auxin reporter R2D2 fluorescence after UV irradiation as shown in B. Nuclear signal intensity in channels for DII-Venus and mDII-ntTomato was measured from cotyledon areas UV-treated (UV) and non-UV-treated (adjacent and distal). n = 28 cells per zone from 4 cotyledons. Representative results from 4 experimental replicates. Plot shows mean (dots) + SEM (dashed lines). (D) Schematic representation of auxin uncaging experiment to investigate the induction of pavement cell interdigitation by uncaged auxin in the region outside of the uncaging site (red square box). UV light indicates the site of uncaging (red oval). This experiment was conducted in 3.5-day-old seedlings overexpressing ARR20-OX to suppress the production of endogenous auxin. (E) Pavement cell phenotypes outside of the UV-treated area, as indicated by a red square box in D, were imaged. Scale bar = 50 µm. (F) Quantitative analysis of pavement cell phenotype shown in E. Violin plot of lobe number per cell (*Left*) and margin roughness (*Right*). Box plot inside each violin plot depicts four quartiles and the median. Red dot depicts the mean value. Raw images were auto segmented and analyzed with PaCeQuant. Eight different cotyledons, each from different seedlings, were analyzed in each treatment. n = 157 cells in mock, n = 187 cells in auxin uncaged. Similar results were obtained in 3 experimental replicates. *t*-test, ****p<0.0001.

We then performed auxin uncaging near the central portion of cotyledons overexpressing *ARR20* (ARR20-OX), which suppressed the initial auxin accumulation at the tip and PC interdigitation (**Figure 2D**). The concentration of caged auxin used for these assays was in the micromolar range as older seedlings (3.5 DAP) already formed the epidermal cuticle and were much less permeable to DMPNB-NAA. This ectopically induced auxin maximum promoted the PC interdigitation in ARR20-OX cotyledons in the region contiguous with the uncaging (**Figure 2D, E**). The effects were detectable locally (within the UV-treated region) as early as 15 h after uncaging (**Figure S2G**) and globally (outside the UV-treated region) at 40 h after uncaging (**Figure 2D, E**). Cotyledons treated with locally uncaged auxin showed increased lobe number per cell and increased margin roughness (**Figure 2F**). These results indicate that a local auxin maximum promotes PC interdigitation in the entire cotyledon epidermis, supporting our hypothesis that a tip auxin maximum globally coordinates PC morphogenesis in cotyledons.

### TIR1/AFB-based nuclear pathway generates the auxin signal for the activation of PC interdigitation

Given the self-organizing nature of auxin ^5^, we speculated that TIR1/AFB-based transcriptional auxin signaling might be involved in the generation of the tip auxin maximum in cotyledons. We analyzed PC phenotype in cotyledons of the quintuple loss-of-function mutant *tir1-1 afb1-3 afb2-3 afb4-8 afb5-5* (*tir1Qt*) ^39^ because *AFB3* is the only member of the nuclear auxin receptor family with very low expression in the cotyledon epidermis (**Figure S3A**). Like previous findings in the *tir1afb123* quadruple mutant (*tir1Qm*) ^40^, siblings from a homozygous *tir1Qt* line displayed variable seedling phenotypes, which we grouped into five classes (**Figure S3B**). Among them, class III (12%) persistently showed a distinctive aborted root and reduced PC interdigitation **(Figure S3B**). *tir1Qt* III cotyledons showed PC interdigitation defects similar to those in *tmk1234 (tmkQ)* (**Figure 3A, S3C)**, a previously reported defect ^10^, which was partially rescued with TMK1-GFP (**Figure S3D**). Interestingly, *tir1Qt* III cotyledons responded to treatment with 20 nM NAA by increasing their lobe number per cell and margin roughness, same as in wild-type cotyledons (**Figure 3A, B and S3E**). The auxin responsiveness in the *tir1Qt* auxin receptor mutant is in sharp contrast to the *tmk1234* (*tmkQ)* mutant, which is fully insensitive to auxin-induced PC interdigitation (**Figure 3A, B and S3E**).

**Figure 3.**
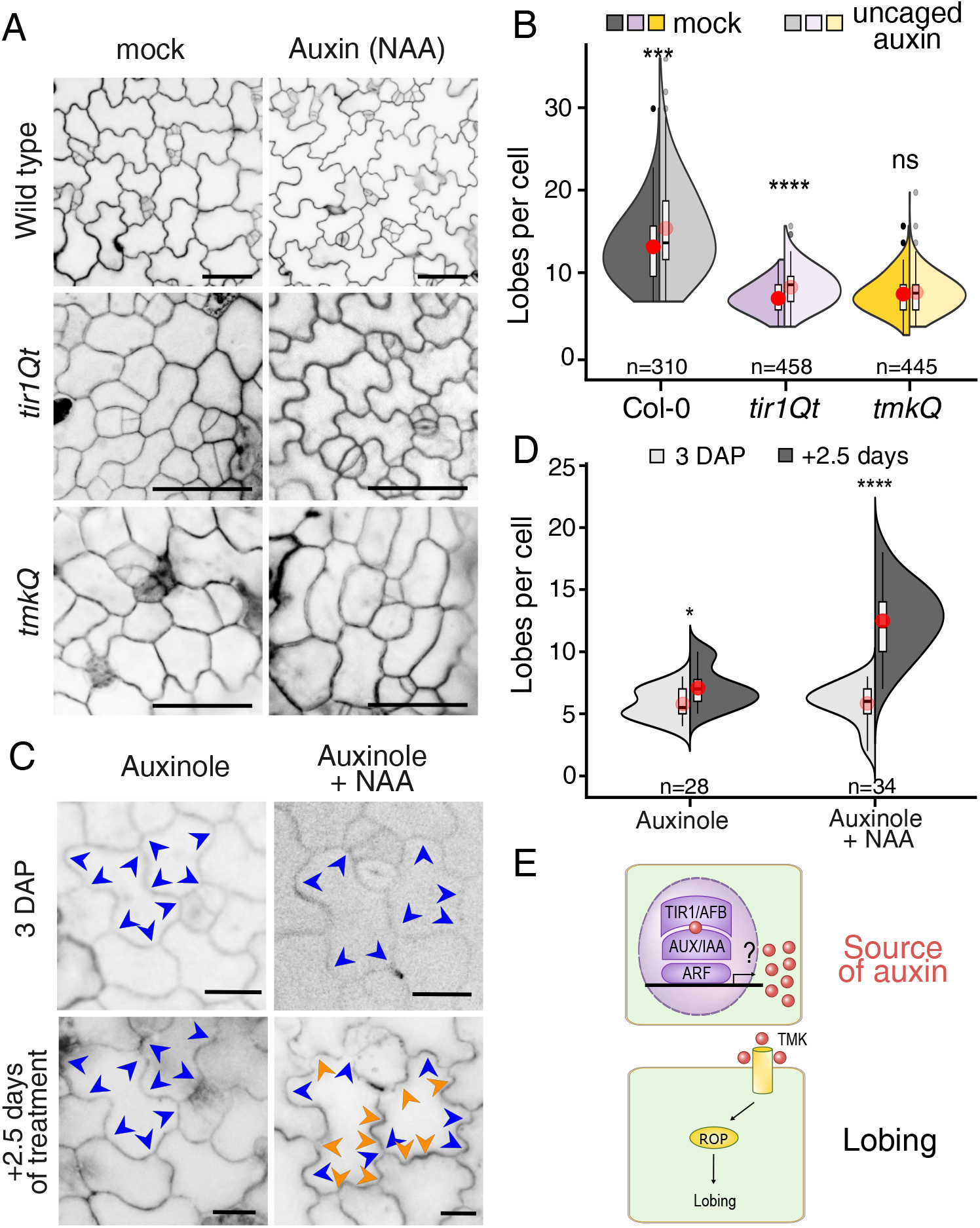
Auxin-induced PC interdigitation in the absence of TIR1/AFBs-based auxin signaling. (A) Exogenous auxin rescues PC interdigitation defects in *tir1Qt* but not in *tmkQ*. Shown are representative images of pavement cells from Col-0, *tir1Qt,* and *tmkQ* seedlings cultured in liquid media with either 0.01% DMSO (mock) or 20 nM auxin NAA (auxin) for 5 days after planting seeds. Scale bar = 50 µm. (B) Quantification of lobe number per cell from images in A. Split violins for each genotype show values obtained from mock (opaque) and NAA-treated (translucent) cotyledons. Box plot inside each violin plot depicts four quartiles and the median. Red dot depicts the average. n is indicated below each plot, with data from at least 8 different cotyledons, each from a different seedling. Similar results were obtained in 5 independent experiments. *t*-test, ns = non-significance, ****p<0.0001. (C) Single-cell tracking experiment showing exogenous auxin-induced lobing in *tir1Qt* seedlings treated with Auxinole. Cotyledons from 3-day-old *tir1Qt* seedlings with existing lobes (blue arrowheads) were treated with 20 µM for 0.5 h before being transferred to a new liquid medium with either 20 µM Auxinole or 20 µM Auxinole +100 nM auxin NAA. The same cells were imaged at the time of mock or NAA treatment and 2.5 days later. Auxin-induced new lobes are indicated with orange arrowheads. (D) Quantitative analysis of PC interdigitation for the single cell tracking experiment described in D. Shown is lobe number per cell before (3 days after plating (DAP), light gray) and after treatment (+2.5 days, dark gray). *t*-test, *p<0.05, ****p<0.0001. (E) Schematic view of the hierarchical auxin system where TIR1/AFBs-dependent auxin synthesis acts as the source for the auxin perceived by TMK-dependent cell-surface auxin signaling.

The responsiveness of *tir1Qt* to auxin could be due to the possible residual signaling activity from AFB3. To test this, we treated the *tir1Qt* mutant with auxinole, which interferes with the TIR1/AFB-dependent degradation of AUX/IAA ^38^. Treatment of wild-type seedlings with auxinole fully reproduced *tir1Qt* class III PC phenotypes (**Figure S4A**) and completely blocked TIR1/AFB-dependent auxin responsiveness measured by histochemical assay using DR5::GUS and by fluorescence using DR5v2 reporter (**Figure S4B**). However, *tir1Qt* cotyledons treated with auxinole remained responsive to NAA-induced PC interdigitation (**Figure S4C).** Notably, *tir1Qt* cotyledons displayed an absence of interdigitation gradient along the proximodistal axis (**Figure S4D**), implying the importance of this auxin signaling pathway in the global coordination. Moreover, single-cell tracking showed that 100 nM NAA induced the formation of new lobes and increased margin roughness in *tir1Qt* PCs even in the presence of auxinole (**Figure 3C, D**). Taken together, our results indicate that TIR1/AFB-based signaling leads to the generation of an auxin signal that directly activates the TMK-dependent PC interdigitation (**Figure 3E**).

The above results with auxinole treatment also suggest that TIR1/AFB-based signaling in PC interdigitation acts to regulate nuclear gene expression, independent of the non-transcriptional function of TIR1/AFBs ^41^. We further tested this by analyzing PC phenotypes in mutants affecting other nuclear components of the TIR1/AFB signaling pathway. By screening available mutations that stabilize Aux/IAA proteins, which repress TIR1/AFB-induced gene expression, we found that *iaa18^D^* (G99E) ^42^ showed very strong defects in PC shape formation compared to wild-type seedlings (**Figure 4A and S4E**). Similar to *tir1Qt*, the defects of PC interdigitation in *iaa18^D^* were restored by exogenously applied auxin (**Figure 4B**). This further corroborates the importance of TIR1/AFB-based transcriptional auxin signaling in promoting PC interdigitation.

**Figure 4.**
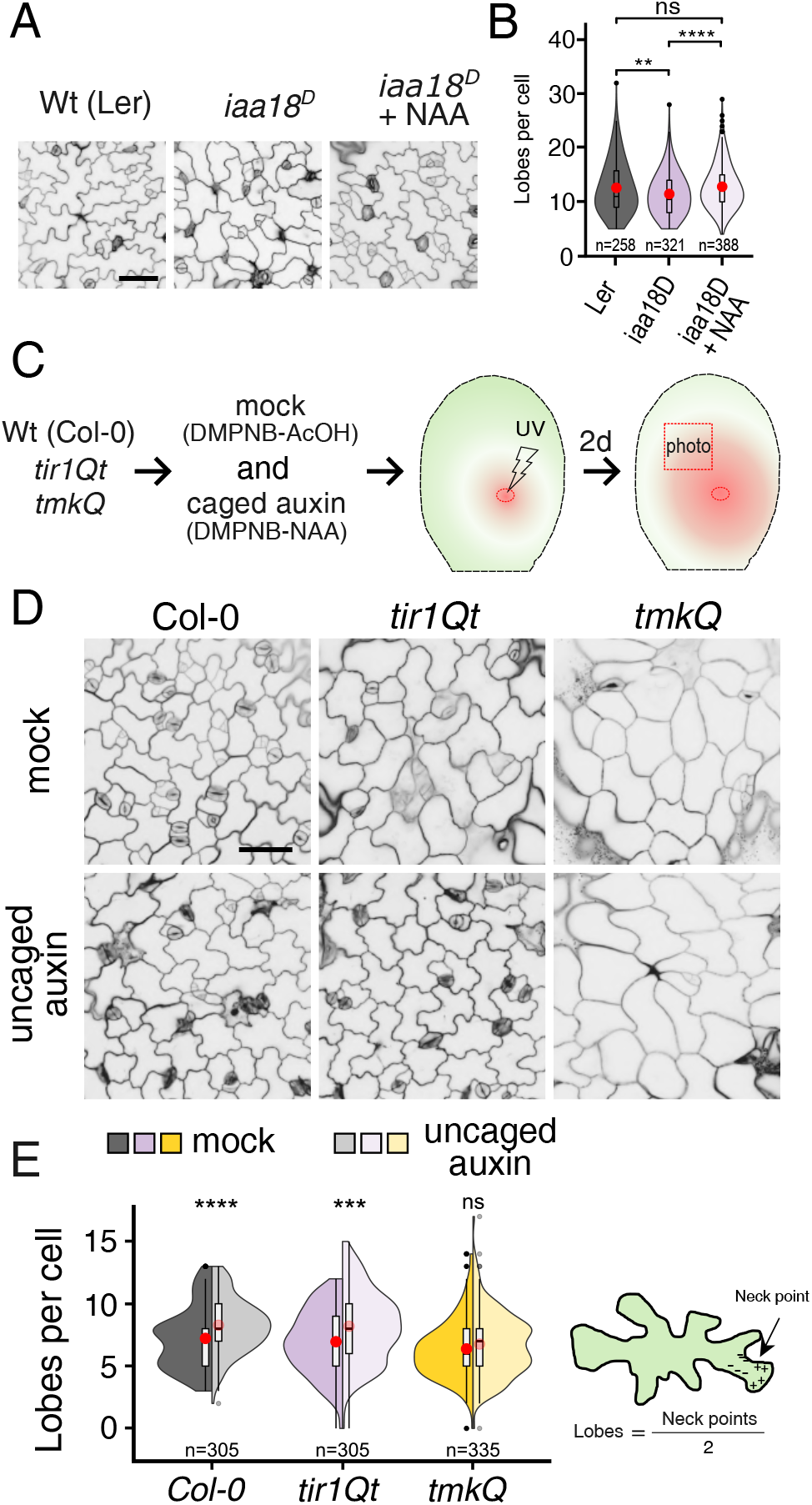
Local auxin uncaging globally rescues defects in pavement cell interdigitation resulting from disruption of the TIR1/AFB signaling pathway. (A) Exogenous auxin treatments restore defects in PC interdigitation observed in *iaa18^D^*. The gain-of-function mutant *iaa18^D^* results from a point mutation in domain II of AUX/IAA protein causing their stabilization and inhibition of auxin transcriptional responses. Seedlings were grown in the absence or presence of 20 nM NAA for 4 days. Scale bars = 50 μm. (B) Quantitative analysis of the PC interdigitation phenotype in *iaa18^D^* mutant as shown in panel B. Statistical analysis showed that the mean lobe number per cell in wild-type cotyledon PCs was significantly greater than in *iaa18^D^* PCs (purple opaque) but not different from *iaa18^D^* PCs treated with NAA (purple translucid). n=250-388 cells from 8 different cotyledons, each from different seedlings. Results representative from 4 experimental replicates. (C) Schematics of the local auxin uncaging protocol. 3.5-days-old seedlings were soaked in either caged-mock (100 <u>μ</u>M DMPNB-AcOH) or caged auxin (100 <u>μ</u>M DMPNB-NAA) for 5 h. Then, seedlings were UV-treated for 30 sec (25% laser, 60 mW) and placed back in semi-solid medium to grow for 2 days. Cotyledons were then excised and stained to analyze cell shape outside of the UV-treated area. (D) Local auxin uncaging globally induced lobing in *tir1Qt* but not in *tmkQ* mutants. Representative images from seedlings treated as shown in C. Scale bar = 50 µm. (E) Quantitative analysis of PC interdigitation as shown in D. Lobe number per cell is shown. For each genotype, split violins show the mock (opaque) and NAA treatment (translucent) values. Box plot inside each violin plot depicts four quartiles and the median. Red dot depicts the average. n = 305-335 cells from 8 different cotyledons, each from different seedlings. Similar results were obtained in 5 independent experiments. *t*-test, ***p<0.001, ****p<0.001.

Finally, we asked if local auxin could overcome defects in TIR1/AFB-dependent global coordination of PC interdigitation, as we would expect if TIR1/AFB-dependent nuclear auxin signaling is needed to generate the tip-high auxin maximum. Thus, we locally increased auxin in *Col-0* wild type and *tirQt* III cotyledons (**Figure 4C**), and in both genotypes observed that local uncaging of auxin significantly increased the lobe number per cell within and outside of the uncaging region (**Figure 4D, E**). This indicates a global response to local uncaging of auxin in the *tir1Qt*, but not in *tmkQ* cotyledons (**Figure 4D, E**). Altogether, our results suggest that the TIR1/AFB-based nuclear pathway generates an apical auxin maximum that acts globally in promoting PC interdigitation in Arabidopsis cotyledons.

### The TIR1/AFB pathway activates the expression of auxin-biosynthetic genes

We next investigated how TIR1/AFB-based transcriptional signaling activates the formation of tip-localized auxin maximum in cotyledons. Mutations in *YUC* or *IBR* genes, which are involved in the YUC-TAA and indole-3-butyric acid (IBA) auxin biosynthesis pathways ^20,43^, respectively, cause defects in PC interdigitation. We examined whether the expression of these auxin-biosynthetic genes was affected in the *tir1Qt* mutant. Quantitative PCR analysis showed that mRNA levels for ECH2 and IBR10, which are functionally redundant and crucial for the IBA-to-IAA conversion pathway ^44^, were greatly reduced in *tir1Qt* young cotyledons (**Figure 5A, S5A, B**). Furthermore, auxin-induced ECH2 and IBR10 gene expression was detected in Col-0 wild type but not in *tir1Qt* (**Figure S5B, C**). Consistently, the double mutant *ech2-/-;ibr10-/-* shows a strong defect in PC interdigitation. Notably, this strong PC phenotype can be rescued with either YFP-IBR10 or YFP-ECH2 (**Figure S5D, E**). Furthermore, *DR5::GUS* expression at the tip of cotyledons was essentially eliminated in the *ech2-/-;ibr10-/-* double mutant, consistently with their PC phenotype (**Figure 5B**). Finally, exogenous auxin fully restores the PC interdigitation defect in the *ech2-/-;ibr10-/-* mutant (**Figure 5C-D**). These results indicate that the IBA-dependent auxin biosynthetic pathway is regulated by the TIR1/AFB-based transcriptional signaling and contributes to the tip-high auxin maximum.

**Figure 5.**
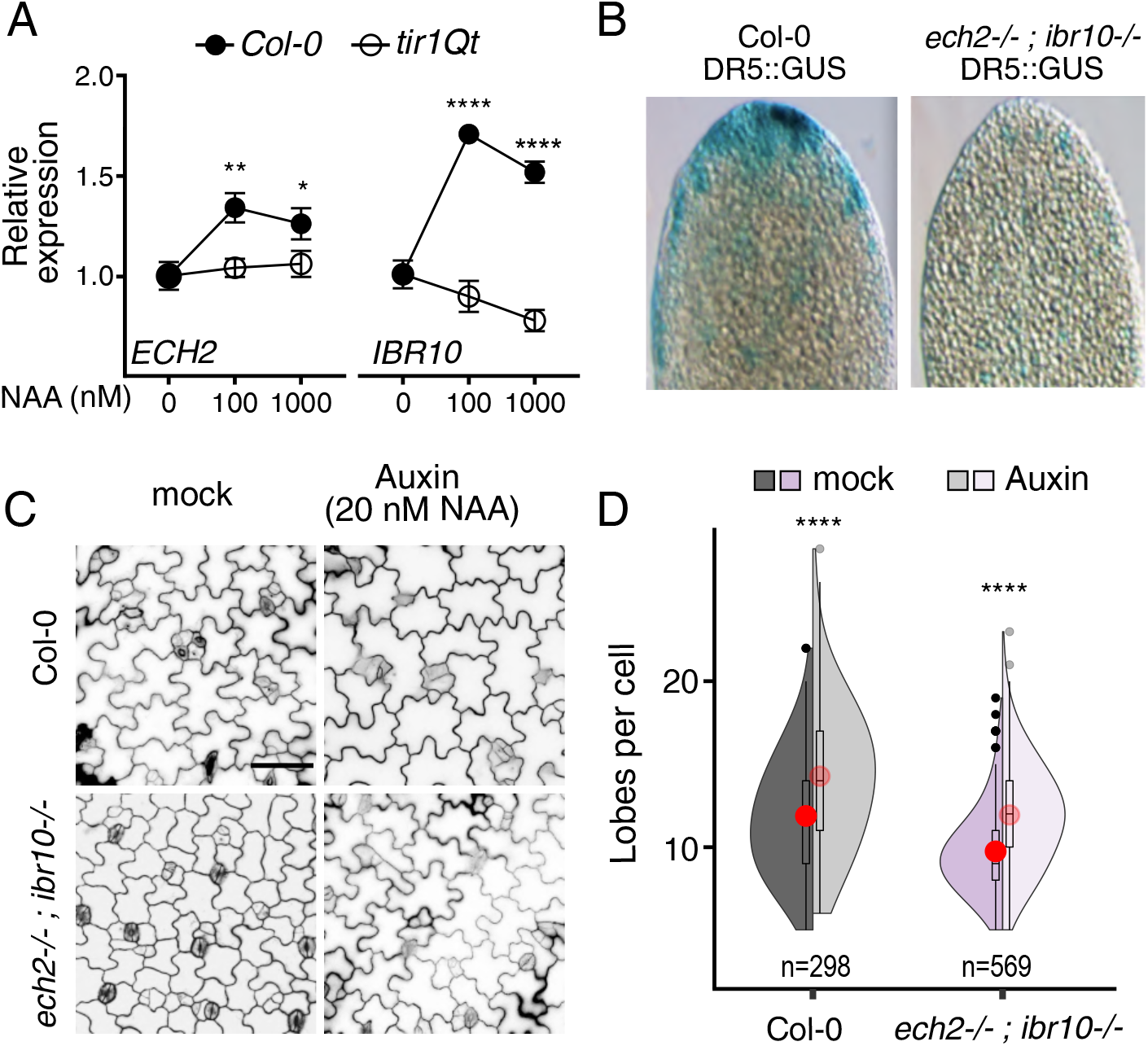
The TIR1/AFB-based nuclear pathway is required for the expression of the IBR auxin-biosynthetic genes that contribute to auxin maxima at the tip of cotyledons. (A) Induction of *ECH2* and *IBR10* gene expression by auxin was compromised in *tir1Qt*. Auxin treatment and qRT-PCR analysis of *ECH2* and *IBR10* expression in wild type Col-0 and *tir1Qt* III as described in Methods. The graph informs 3 biological replicates, each reaction is performed with 3 technical replicates. t-test, *p<0.05. (B) Tip-high *DR5::GUS* expression in 48 HAP cotyledons was greatly reduced in the *ech2-/-;ibr10-/-* double mutant. (C) Auxin restored the PC interdigitation defect in the *ech2-/-;ibr10-/-* mutant. Seedlings were grown for 4 days in 20 nM NAA. Scale bar = 50 µm. (D) Lobes per cell of cotyledons shown in C. Split violins show mock (opaque) and NAA treatment (translucent) values, for each genotype. Box plot inside each violin plot depicts four quartiles and the median. Red dot depicts the average. Split violins for each genotype show values obtained from mock (opaque) and NAA-treated (translucent) cotyledons. n = 298-569 cells from 10 cotyledons. *t*-test, ****p<0.0001.

### TIR1/AFB-dependent auxin signal locally activates ROP signaling and PC interdigitation

The auxin signal generated by the TIR1/AFB pathway may directly activate TMK-dependent ROP2 and ROP6 to establish PC interdigitation or may promote cell expansion resulting in mechanical stress, which has been proposed to activate PC interdigitation ^45,46^. As a first step in distinguishing these two possible models, we locally induce an auxin maximum and evaluated PC phenotypes. PCs from *Col-0* and *tir1Qt* cotyledons responded to locally uncaged auxin, however, local auxin did not activate PC morphogenesis in *tmkQ* as measured by the lobe number per cell (**Figure 4D, E)**. These results strongly suggest that TMK-based auxin perception and signaling is required for local auxin-induced establishment of PC interdigitation, as previously reported ^10,12^. Thus, we propose that the TIR1/AFB-dependent auxin signal, once reaching a specific cell, will locally activate the interdigitation of that specific cell. To further test this hypothesis, we examined changes in auxin-induced ROP2 and ROP6 activity after a prolonged auxinole treatments of wild-type *Col-0* seedlings to eliminate the TIR1/AFB-based auxin signaling. We found that ROP2 and ROP6 activity was reduced by auxinole treatments, and that this reduction was reversed by exogenous auxin. More importantly, exogenous auxin activates ROP2 and ROP6 activity equally in mock as in auxinole treatments (**Figure S6A, B**). Consistently, *rop2;rop4;rop6* mutants remain insensitive to exogenous auxin treatments in promoting lobe formation (**Figure S6E, F**). Altogether these results suggest that an upstream TIR1/AFBs-based nuclear auxin signaling pathway generates an auxin signal that locally activates PC interdigitation directly through the TMK-dependent ROP signaling pathways.

### Auxin establishes PC interdigitation decoupled from cell expansion-induced associated mechanical stress

We further investigated whether auxin directly activates cell polarization pathways or promotes PC interdigitation indirectly through cell expansion-derived mechanical stress. In the latter case, cell expansion, either dependent or independent of auxin, is expected to promote PC interdigitation. Thus, we monitored the birth of the interdigitation period (0 - 48 HAP) along the proximo-distal axis of cotyledons (**Figure S6C).** We extracted three shape metrics: cell area, largest empty circle (LEC), and margin roughness (MR). LEC serves as a proxy for mechanical stress magnitude experienced by individual cells and is proposed to be low in cells with complex shapes ^47,48^. Local LEC (LLEC) ^48^ only differs from LEC once curvature is formed, thus, not appropriate for this analysis. Meanwhile, MR is a proxy for interdigitation status by measuring local curvature around the border ^24^. If interdigitation is established from cell expansion-induced mechanical stress, LEC would be expected to correlate with MR in expanding cells. Interestingly, we found no correlation during 0-48 HAP between MR and LEC (R^2^ <0.02) (**Figure 6A, upper panel**). Additionally, if mechanical stress were to activate PC interdigitation, greater expansion in expanding cells would be tightly linked with greater interdigitation. However, we found a very weak correlation between interdigitation and cell size or area (R^2^ <0.16) (**Figure 6A, lower panel**) in the 0-48 HAP period. Most of that positive correlation was contributed by cells at the tip in 48 HAP cotyledons coinciding with the auxin maximum (**Figure S6D**). These results suggest that an increase in cell size and mechanical stress does not necessarily promote interdigitation.

**Figure 6.**
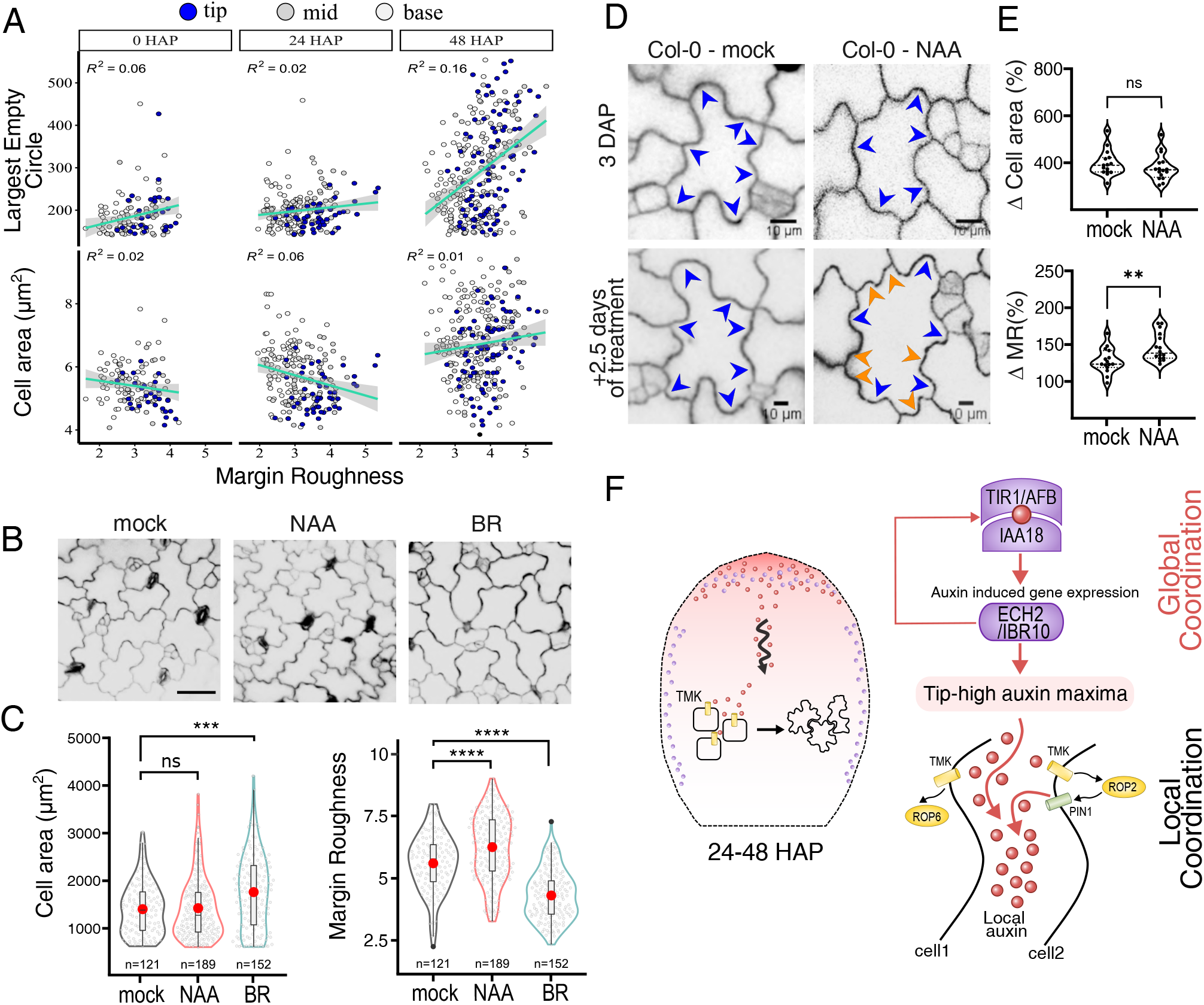
Auxin-induced PC interdigitation is decoupled from cell expansion-induced mechanical stress. (A) PC interdigitation does not correlate with mechanical stress or cell size in early-developing cotyledons. Shown are correlation plots between margin roughness (MR) and the largest empty circle (LEC, upper graph), which is indicative of the mechanical stress ^47^ and between MR and cell size/area (lower panel) at 0 HAP, 24 HAP and 48 HAP and at different positions (tip, base, middle) in the cotyledon. Green line is the linear model. Gray shadows display the 95% confidence interval. R^2^ = correlation coefficient. (B) Cell expansion without PC interdigitation. 24 HAP wild-type seedlings were either mock-treated or treated with 1 µM auxin NAA or 1 µM brassinolide for 4 days. Then, cotyledons were stained and imaged by confocal microscopy for posterior analysis with PaCeQuant. Growth for 1 day before treatment is crucial to avoid auxin-induced inhibition of germination. (C) Violin plot of cell size (left) and margin roughness (right) computed from images as shown in B. n > 51-109 cells from 9 cotyledons. Box plot inside each violin plot depicts four quartiles and the median. Red dot depicts the mean value. Wilcox test, *p<0.05, ****p<0.0001. (D) Auxin-induced *de novo* lobe formation without increasing cell size in single cell tracking experiments. Cotyledons (3 DAP) with formed lobes (blue arrowheads) were mock-treated (diluted DMSO) or treated with 20 nM auxin NAA for 2.5 days and analyzed as described in Figure 3C. (E) Percentage variation (Δ) in cell size (top) and margin roughness (bottom) calculated with pre/post treatment pairwise images. n = 15 cells, from 5 cotyledons each from different seedlings. Same results were obtained in 3 independent experiments. *t*-test, ** p<0.01, ns = non-significance. (F) A model for a hierarchical global and local auxin signaling systems underlying the PC interdigitation pattern. A basal level of auxin, which self-amplifies via TIR1/AFB1-dependent auxin signaling to activate IBR-dependent auxin synthesis genes (purple dots). This is counteracted by cytokinin signaling, restricting auxin maxima (increased red color) to the tip of cotyledons. The auxin maxima act as a global signal by emanating to the remaining regions of the cotyledon epidermis (wavy black arrow) presumably via diffusion through the apoplastic space, which locally increases the level of auxin for a specific cell. The resultant local auxin (red dots) then triggers TMK/ROP-dependent cell polarization and cell-cell coordination by activating the feedback loop and the complementary ROP2/ROP6 pathways to coordinate lobe and indentation formation ^10,16,20,21^.

To further assess whether cell expansion is the cause of PC interdigitation, we synchronously increased cell sizes by treating cotyledons with 1 μM brassinosteroids (BR). This treatment increased cell size by 50% but decreased MR by 40% (**Figure 6B, C**), agreeing with a recent report ^49^. In contrast, NAA treatments greatly increased MR without increasing cell sizes (**Figure 6B, C).** Thus, we conclude that cell expansion-induced mechanical stress is unlikely the driving force for PC interdigitation induced by auxin. Finally, tracking of mock- or NAA-treated PCs with already consolidated interdigitation (**Figure 6D**, blue arrows) revealed that cells in both conditions exhibited a 4-fold increase in cell size in 2 days (**Figure 6D**), but only NAA-treated cells increased MR by 50% (**Figure 6E**). More importantly, only NAA-treated cells displayed new lobes after treatment (**Figure 6D**, orange arrows). Altogether, our results suggest that local auxin promotes PC interdigitation directly by activating the TMK-dependent formation of PC multi-polarity and not indirectly via auxin-induced cell expansion and the resulting mechanical stress.

### Conclusions and Discussion

Here we show that two auxin signaling systems, a TIR1/AFB-based nuclear signaling and a TMK-based cell surface signaling, coordinately control PC interdigitation in Arabidopsis cotyledons. Our findings suggest that they act at different functional scales and in a hierarchical manner (**Figure 6F).** At the whole organ level, TIR1/AFBs-based transcriptional signaling amplifies the initial auxin signal in part via activating the expression of auxin biosynthetic genes, leading to the generation of the auxin maximum at the tip of cotyledons (**Figure 6F, red gradient**). As auxin moves across the entire surface of the cotyledon from tip to base, auxin locally activates PC interdigitation via TMK-based cell surface signaling and ROP activation ^10,12^. This hierarchical relationship between the two auxin signaling mechanisms integrates global coordination with local activation of PC interdigitation throughout the entire epidermis.

PC interdigitation is globally coordinated by TIR1/AFBs-based nuclear auxin signaling that is restricted to the cotyledon tip and margins by repressive cytokinin signaling. This global signaling is a self-organizing process relying on the TIR1/AFB-activated transcription of genes involved in the synthesis of IBA-derived auxin (**Figure 6E, purple dots**), which together with base-to-tip auxin transport along the cotyledon’s margins generates a transient tip-high auxin maximum ^50^. Time-lapse imaging shows that auxin at the cotyledon tip rapidly propagates to the rest of the cotyledon, but the mode of this propagation remains to be determined. The tip-derived auxin appears to override local auxin gradients observed around stomata cells ^51^, because *spch* mutants, which lack the stomata cell lineage, display the same tip-to-base interdigitation gradient observed in wild type ^52^.

PC interdigitation is locally coordinated by a TMK-based auxin signaling module for lobe formation (**Figures 3 and 4**). This local activation of PC interdigitation is also a self-organizing process. ROP2-dependent polarization of the PIN1 auxin efflux carrier generates local extracellular auxin that coordinately activates ROP2 and ROP6 between neighboring cells ^20,23^. Thus, this local auxin signaling mechanism generates differential features along the PC contour, such as differential pectin accumulation and differential cell wall strength, which is accompanied by and may be reinforced by mechanical signals (**Figure 6**) ^45,53–56^.

The hierarchical self-organizing morphogenetic mechanism we reveal here for the Arabidopsis cotyledon may also be controlling planar polarity in roots ^7,8^ and is analogous to WNT signaling regulating planar cell polarity (PCP) in animal systems. Similar to TIR1/AFB nuclear auxin signaling, the canonical WNT11 signaling pathway activates the transcription of genes proposed to instruct global coordination of PCP-mediated processes such as body axis formation and orientation of hairs ^1,57,58^. Similar to the TMK-ROP signaling, WNT11 also activates the Rho GTP-dependent pathway, which locally coordinates PCP establishment that is required for myocyte orientation and elongation of embryonic muscle fibers ^59^. Therefore, the control of these developmental processes in plants and animals appears to share general design principles, although the details of the molecular mechanisms are quite different.

It remains to be seen whether such hierarchically coordinated self-organizing auxin signaling systems also regulate other developmental and morphogenetic processes in plants. Nonetheless, the TIR1/AFB-based nuclear auxin signaling and the TMK-based cell surface auxin signaling appear to coordinately regulate other auxin-dependent processes such as pH-mediated hypocotyl elongation, root growth, and lateral root formation ^13,15,60–62^. Furthermore, auxin regulates the polarization of PIN proteins and the activity of auxin biosynthetic through these two different pathways ^14,63^. Hence investigating the biological significance and the mechanisms behind the coordination between these two distinct auxin signaling pathways will be an exciting and fertile field of inquiry in the years to come.

### Limitations to the study

As discussed above, our study described here strongly indicates that that the TIR1/AFB pathway underlies the global coordination of pavement cell morphogenesis in Arabidopsis cotyledons. However, our study does have a limitation in that the siblings of the *tir1Qt* mutant exhibit highly variable phenotypes, making it extremely difficult to perform clonal analysis that would provide additional support for this conclusion. This limitation also hinders a genetic experiment that could further test the functional relationship between the TIR1/AFB and TMK pathways. Additionally, our study did not address the mechanistic details of the auxin dynamics in the cotyledon. In this work, we propose a tip-to-base apoplastic auxin diffusion; however, auxin movement mediated by auxin transports could not be excluded.

## Acknowledgments

We are grateful to Natasha Raikhel for the helpful suggestions. This work is supported in part by grants from the U.S. National Institute of General Medical Sciences (GM081451 and GM100130) and Shenzhen University of Advanced Technology startup funds to ZY and the National Key Laboratory of Quantitative Synthetic Biology, Shenzhen Institutes of Advanced Technology, Chinese Academy of Sciences, from the European Research Council (project ERC-2011-StG-20101109-PSDP) and CEITEC – Central European Institute of Technology (CZ.1.05/ 1.1.00/02.0068) to J.F.

## References

1. Aw, W.Y., and Devenport, D. (2017). Planar cell polarity: global inputs establishing cellular asymmetry. Curr. Opin. Cell Biol. 10.1016/j.ceb.2016.08.002.

2. Torban, E., and Sokol, S.Y. (2021). Planar cell polarity pathway in kidney development, function and disease. Nat. Rev. Nephrol. 17, 369–385. 10.1038/s41581-021-00395-6.

3. Butler, M.T., and Wallingford, J.B. (2017). Planar cell polarity in development and disease. Nat. Rev. Mol. Cell Biol. 18, 375–388. 10.1038/nrm.2017.11.

4. Angers, S., and Moon, R.T. (2009). Proximal events in Wnt signal transduction. Nat. Rev. Mol. Cell Biol. 10, 468–477. 10.1038/nrm2717.

5. Friml, J. (2022). Fourteen stations of auxin. Cold Spring Harb. Perspect. Biol. 14. 10.1101/cshperspect.a039859.

6. Hammes, U.Z., Murphy, A.S., and Schwechheimer, C. (2021). Auxin Transporters-A Biochemical View. Cold Spring Harb. Perspect. Biol. 10.1101/cshperspect.a039875.

7. Ikeda, Y., Men, S., Fischer, U., Stepanova, A.N., Alonso, J.M., Ljung, K., and Grebe, M. (2009). Local auxin biosynthesis modulates gradient-directed planar polarity in Arabidopsis. Nat. Cell Biol. 11, 731–738. 10.1038/ncb1879.

8. Fischer, U., Ikeda, Y., Ljung, K., Serralbo, O., Singh, M., Heidstra, R., Palme, K., Scheres, B., and Grebe, M. (2006). Vectorial information for Arabidopsis planar polarity is mediated by combined AUX1, EIN2, and GNOM activity. Curr. Biol. 16, 2143–2149. 10.1016/j.cub.2006.08.091.

9. Weijers, D., and Wagner, D. (2016). Transcriptional responses to the auxin hormone. Annu. Rev. Plant Biol. 67, 539–574. 10.1146/annurev-arplant-043015-112122.

10. Xu, T., Dai, N., Chen, J., Nagawa, S., Cao, M., Li, H., Zhou, Z., Chen, X., De Rycke, R., Rakusová, H., et al. (2014). Cell surface ABP1-TMK auxin-sensing complex activates ROP GTPase signaling. Science 343, 1025–1028. 10.1126/science.1245125.

11. Cao, M., Chen, R., Li, P., Yu, Y., Zheng, R., Ge, D., Zheng, W., Wang, X., Gu, Y., Gelová, Z., et al. (2019). TMK1-mediated auxin signalling regulates differential growth of the apical hook. Nature 568, 240–243. 10.1038/s41586-019-1069-7.

12. Pan, X., Fang, L., Liu, J., Senay-Aras, B., Lin, W., Zheng, S., Zhang, T., Guo, J., Manor, U., Van Norman, J., et al. (2020). Auxin-induced signaling protein nanoclustering contributes to cell polarity formation. Nat. Commun. 11, 3914. 10.1038/s41467-020-17602-w.

13. Huang, R., Zheng, R., He, J., Zhou, Z., Wang, J., Xiong, Y., and Xu, T. (2019). Noncanonical auxin signaling regulates cell division pattern during lateral root development. Proc Natl Acad Sci USA 116, 21285–21290. 10.1073/pnas.1910916116.

14. Wang, Q., Qin, G., Cao, M., Chen, R., He, Y., Yang, L., Zeng, Z., Yu, Y., Gu, Y., Xing, W., et al. (2020). A phosphorylation-based switch controls TAA1-mediated auxin biosynthesis in plants. Nat. Commun. 11, 679. 10.1038/s41467-020-14395-w.

15. Lin, W., Zhou, X., Tang, W., Takahashi, K., Pan, X., Dai, J., Ren, H., Zhu, X., Pan, S., Zheng, H., et al. (2021). TMK-based cell-surface auxin signalling activates cell-wall acidification. Nature 599, 278–282. 10.1038/s41586-021-03976-4.

16. Yu, Y., Tang, W., Lin, W., Li, W., Zhou, X., Li, Y., Chen, R., Zheng, R., Qin, G., Cao, W., et al. (2023). ABLs and TMKs are co-receptors for extracellular auxin. Cell 186, 5457–5471.e17. 10.1016/j.cell.2023.10.017.

17. Pérez-Henríquez, P., and Yang, Z. (2023). Extranuclear auxin signaling: a new insight into auxin’s versatility. New Phytol. 237, 1115–1121. 10.1111/nph.18602.

18. Liu, S., Jobert, F., Rahneshan, Z., Doyle, S.M., and Robert, S. (2021). Solving the puzzle of shape regulation in plant epidermal pavement cells. Annu. Rev. Plant Biol. 72, 525–550. 10.1146/annurev-arplant-080720-081920.

19. Lin, W., and Yang, Z. (2020). Unlocking the mechanisms behind the formation of interlocking pavement cells. Curr. Opin. Plant Biol. 57, 142–154. 10.1016/j.pbi.2020.09.002.

20. Xu, T., Wen, M., Nagawa, S., Fu, Y., Chen, J.-G., Wu, M.-J., Perrot-Rechenmann, C., Friml, J., Jones, A.M., and Yang, Z. (2010). Cell surface- and rho GTPase-based auxin signaling controls cellular interdigitation in Arabidopsis. Cell 143, 99–110. 10.1016/j.cell.2010.09.003.

21. Fu, Y., Gu, Y., Zheng, Z., Wasteneys, G., and Yang, Z. (2005). Arabidopsis interdigitating cell growth requires two antagonistic pathways with opposing action on cell morphogenesis. Cell 120, 687–700. 10.1016/j.cell.2004.12.026.

22. Fu, Y., Xu, T., Zhu, L., Wen, M., and Yang, Z. (2009). A ROP GTPase signaling pathway controls cortical microtubule ordering and cell expansion in Arabidopsis. Curr. Biol. 19, 1827–1832. 10.1016/j.cub.2009.08.052.

23. Nagawa, S., Xu, T., Lin, D., Dhonukshe, P., Zhang, X., Friml, J., Scheres, B., Fu, Y., and Yang, Z. (2012). ROP GTPase-dependent actin microfilaments promote PIN1 polarization by localized inhibition of clathrin-dependent endocytosis. PLoS Biol. 10, e1001299. 10.1371/journal.pbio.1001299.

24. Möller, B., Poeschl, Y., Plötner, R., and Bürstenbinder, K. (2017). PaCeQuant: A Tool for High-Throughput Quantification of Pavement Cell Shape Characteristics. Plant Physiol. 175, 998–1017. 10.1104/pp.17.00961.

25. Nowak, J., Eng, R.C., Matz, T., Waack, M., Persson, S., Sampathkumar, A., and Nikoloski, Z. (2021). A network-based framework for shape analysis enables accurate characterization of leaf epidermal cells. Nat. Commun. 12, 458. 10.1038/s41467-020-20730-y.

26. Wu, T.-C., Belteton, S.A., Pack, J., Szymanski, D.B., and Umulis, D.M. (2016). LobeFinder: A Convex Hull-Based Method for Quantitative Boundary Analyses of Lobed Plant Cells. Plant Physiol. 171, 2331–2342. 10.1104/pp.15.00972.

27. Benková, E., Michniewicz, M., Sauer, M., Teichmann, T., Seifertová, D., Jürgens, G., and Friml, J. (2003). Local, efflux-dependent auxin gradients as a common module for plant organ formation. Cell 115, 591–602. 10.1016/s0092-8674(03)00924-3.

28. Chen, J.G., Shimomura, S., Sitbon, F., Sandberg, G., and Jones, A.M. (2001). The role of auxin-binding protein 1 in the expansion of tobacco leaf cells. Plant J. 28, 607–617. 10.1046/j.1365-313x.2001.01152.x.

29. Brunoud, G., Wells, D.M., Oliva, M., Larrieu, A., Mirabet, V., Burrow, A.H., Beeckman, T., Kepinski, S., Traas, J., Bennett, M.J., et al. (2012). A novel sensor to map auxin response and distribution at high spatio-temporal resolution. Nature 482, 103–106. 10.1038/nature10791.

30. Dello Ioio, R., Nakamura, K., Moubayidin, L., Perilli, S., Taniguchi, M., Morita, M.T., Aoyama, T., Costantino, P., and Sabatini, S. (2008). A genetic framework for the control of cell division and differentiation in the root meristem. Science 322, 1380–1384. 10.1126/science.1164147.

31. Nakamura, A., Higuchi, K., Goda, H., Fujiwara, M.T., Sawa, S., Koshiba, T., Shimada, Y., and Yoshida, S. (2003). Brassinolide induces IAA5, IAA19, and DR5, a synthetic auxin response element in Arabidopsis, implying a cross talk point of brassinosteroid and auxin signaling. Plant Physiol. 133, 1843–1853. 10.1104/pp.103.030031.

32. Marhavý, P., Duclercq, J., Weller, B., Feraru, E., Bielach, A., Offringa, R., Friml, J., Schwechheimer, C., Murphy, A., and Benková, E. (2014). Cytokinin controls polarity of PIN1-dependent auxin transport during lateral root organogenesis. Curr. Biol. 24, 1031–1037. 10.1016/j.cub.2014.04.002.

33. Müller, B., and Sheen, J. (2008). Cytokinin and auxin interaction in root stem-cell specification during early embryogenesis. Nature 453, 1094–1097. 10.1038/nature06943.

34. Li, H., Xu, T., Lin, D., Wen, M., Xie, M., Duclercq, J., Bielach, A., Kim, J., Reddy, G.V., Zuo, J., et al. (2013). Cytokinin signaling regulates pavement cell morphogenesis in Arabidopsis. Cell Res. 23, 290–299. 10.1038/cr.2012.146.

35. D’Agostino, I.B., Deruère, J., and Kieber, J.J. (2000). Characterization of the response of the Arabidopsis response regulator gene family to cytokinin. Plant Physiol. 124, 1706–1717. 10.1104/pp.124.4.1706.

36. Kusaka, N., Maisch, J., Nick, P., Hayashi, K., and Nozaki, H. (2009). Manipulation of intracellular auxin in a single cell by light with esterase-resistant caged auxins. Chembiochem 10, 2195–2202. 10.1002/cbic.200900289.

37. Hayashi, K.-I., Kusaka, N., Yamasaki, S., Zhao, Y., and Nozaki, H. (2015). Development of 4-methoxy-7-nitroindolinyl (MNI)-caged auxins which are extremely stable in planta. Bioorg. Med. Chem. Lett. 25, 4464–4471. 10.1016/j.bmcl.2015.09.001.

38. Hayashi, K., Neve, J., Hirose, M., Kuboki, A., Shimada, Y., Kepinski, S., and Nozaki, H. (2012). Rational design of an auxin antagonist of the SCF(TIR1) auxin receptor complex. ACS Chem. Biol. 7, 590–598. 10.1021/cb200404c.

39. Prigge, M.J., Platre, M., Kadakia, N., Zhang, Y., Greenham, K., Szutu, W., Pandey, B.K., Bhosale, R.A., Bennett, M.J., Busch, W., et al. (2020). Genetic analysis of the Arabidopsis TIR1/AFB auxin receptors reveals both overlapping and specialized functions. eLife 9. 10.7554/eLife.54740.

40. Parry, G., Calderon-Villalobos, L.I., Prigge, M., Peret, B., Dharmasiri, S., Itoh, H., Lechner, E., Gray, W.M., Bennett, M., and Estelle, M. (2009). Complex regulation of the TIR1/AFB family of auxin receptors. Proc Natl Acad Sci USA 106, 22540–22545. 10.1073/pnas.0911967106.

41. Serre, N.B.C., Kralík, D., Yun, P., Slouka, Z., Shabala, S., and Fendrych, M. (2021). AFB1 controls rapid auxin signalling through membrane depolarization in Arabidopsis thaliana root. Nat. Plants. 10.1038/s41477-021-00969-z.

42. Ploense, S.E., Wu, M.-F., Nagpal, P., and Reed, J.W. (2009). A gain-of-function mutation in IAA18 alters Arabidopsis embryonic apical patterning. Development 136, 1509–1517. 10.1242/dev.025932.

43. Strader, L.C., Wheeler, D.L., Christensen, S.E., Berens, J.C., Cohen, J.D., Rampey, R.A., and Bartel, B. (2011). Multiple Facets of Arabidopsis Seedling Development Require Indole-3-Butyric Acid-Derived Auxin. Plant Cell 23, 984–999.

44. Strader, L.C., Wheeler, D.L., Christensen, S.E., Berens, J.C., Cohen, J.D., Rampey, R.A., and Bartel, B. (2011). Multiple facets of Arabidopsis seedling development require indole-3-butyric acid-derived auxin. Plant Cell 23, 984–999. 10.1105/tpc.111.083071.

45. Bidhendi, A.J., Altartouri, B., Gosselin, F.P., and Geitmann, A. (2019). Mechanical stress initiates and sustains the morphogenesis of wavy leaf epidermal cells. Cell Rep. 28, 1237–1250.e6. 10.1016/j.celrep.2019.07.006.

46. Belteton, S.A., Li, W., Yanagisawa, M., Hatam, F.A., Quinn, M.I., Szymanski, M.K., Marley, M.W., Turner, J.A., and Szymanski, D.B. (2021). Real-time conversion of tissue-scale mechanical forces into an interdigitated growth pattern. Nat. Plants 7, 826–841. 10.1038/s41477-021-00931-z.

47. Sapala, A., Runions, A., Routier-Kierzkowska, A.-L., Das Gupta, M., Hong, L., Hofhuis, H., Verger, S., Mosca, G., Li, C.-B., Hay, A., et al. (2018). Why plants make puzzle cells, and how their shape emerges. eLife 7. 10.7554/eLife.32794.

48. Eng, R.C., Schneider, R., Matz, T.W., Carter, R., Ehrhardt, D.W., Jönsson, H., Nikoloski, Z., and Sampathkumar, A. (2021). KATANIN and CLASP function at different spatial scales to mediate microtubule response to mechanical stress in Arabidopsis cotyledons. Curr. Biol. 31, 3262–3274.e6. 10.1016/j.cub.2021.05.019.

49. Liu, X., Yang, Q., Wang, Y., Wang, L., Fu, Y., and Wang, X. (2018). Brassinosteroids regulate pavement cell growth by mediating BIN2-induced microtubule stabilization. J. Exp. Bot. 69, 1037–1049. 10.1093/jxb/erx467.

50. Pérez-Henríquez, P., Nagawa, S., Liu, Z., Pan, X., Michniewicz, M., Tang, W., Rasmussen, C., Van Norman, J., Strader, L., and Yang, Z. (2022). PIN2-mediated self-organizing transient auxin flow contributes to auxin maxima at the tip of Arabidopsis cotyledons.

51. Grones, P., Majda, M., Doyle, S.M., Van Damme, D., and Robert, S. (2020). Fluctuating auxin response gradients determine pavement cell-shape acquisition. Proc Natl Acad Sci USA 117, 16027–16034. 10.1073/pnas.2007400117.

52. Mansfield, C., Newman, J.L., Olsson, T.S.G., Hartley, M., Chan, J., and Coen, E. (2018). Ectopic BASL Reveals Tissue Cell Polarity throughout Leaf Development in Arabidopsis thaliana. Curr. Biol. 28, 2638–2646.e4. 10.1016/j.cub.2018.06.019.

53. Tang, W., Lin, W., Zhou, X., Guo, J., Dang, X., Li, B., Lin, D., and Yang, Z. (2022). Mechano-transduction via the pectin-FERONIA complex activates ROP6 GTPase signaling in Arabidopsis pavement cell morphogenesis. Curr. Biol. 32, 508–517.e3. 10.1016/j.cub.2021.11.031.

54. Lin, W., Tang, W., Pan, X., Huang, A., Gao, X., Anderson, C.T., and Yang, Z. (2022). Arabidopsis pavement cell morphogenesis requires FERONIA binding to pectin for activation of ROP GTPase signaling. Curr. Biol. 32, 497–507.e4. 10.1016/j.cub.2021.11.030.

55. Majda, M., Grones, P., Sintorn, I.-M., Vain, T., Milani, P., Krupinski, P., Zagórska-Marek, B., Viotti, C., Jönsson, H., Mellerowicz, E.J., et al. (2017). Mechanochemical polarization of contiguous cell walls shapes plant pavement cells. Dev. Cell 43, 290–304.e4. 10.1016/j.devcel.2017.10.017.

56. Altartouri, B., Bidhendi, A.J., Tani, T., Suzuki, J., Conrad, C., Chebli, Y., Liu, N., Karunakaran, C., Scarcelli, G., and Geitmann, A. (2019). Pectin chemistry and cellulose crystallinity govern pavement cell morphogenesis in a multi-step mechanism. Plant Physiol. 181, 127–141. 10.1104/pp.19.00303.

57. Tao, Q., Yokota, C., Puck, H., Kofron, M., Birsoy, B., Yan, D., Asashima, M., Wylie, C.C., Lin, X., and Heasman, J. (2005). Maternal wnt11 activates the canonical wnt signaling pathway required for axis formation in Xenopus embryos. Cell 120, 857–871. 10.1016/j.cell.2005.01.013.

58. Mlodzik, M. (2020). Planar cell polarity: moving from single cells to tissue-scale biology. Development 147. 10.1242/dev.186346.

59. Schlessinger, K., Hall, A., and Tolwinski, N. (2009). Wnt signaling pathways meet Rho GTPases. Genes Dev. 23, 265–277. 10.1101/gad.1760809.

60. Li, L., Verstraeten, I., Roosjen, M., Takahashi, K., Rodriguez, L., Merrin, J., Chen, J., Shabala, L., Smet, W., Ren, H., et al. (2021). Cell surface and intracellular auxin signalling for H+ fluxes in root growth. Nature 599, 273–277. 10.1038/s41586-021-04037-6.

61. Du, M., Bou Daher, F., Liu, Y., Steward, A., Tillmann, M., Zhang, X., Wong, J.H., Ren, H., Cohen, J.D., Li, C., et al. (2022). Biphasic control of cell expansion by auxin coordinates etiolated seedling development. Sci. Adv. 8, eabj1570. 10.1126/sciadv.abj1570.

62. De Smet, I., Tetsumura, T., De Rybel, B., Frei dit Frey, N., Laplaze, L., Casimiro, I., Swarup, R., Naudts, M., Vanneste, S., Audenaert, D., et al. (2007). Auxin-dependent regulation of lateral root positioning in the basal meristem of Arabidopsis. Development 134, 681–690. 10.1242/dev.02753.

63. Hajný, J., Prát, T., Rydza, N., Rodriguez, L., Tan, S., Verstraeten, I., Domjan, D., Mazur, E., Smakowska-Luzan, E., Smet, W., et al. (2020). Receptor kinase module targets PIN-dependent auxin transport during canalization. Science 370, 550–557. 10.1126/science.aba3178.

