## supplementary information for "Hierarchical global and local auxin signals coordinate cellular interdigitation in *Arabidopsis*"

### Materials and Methods

#### Growth conditions

Arabidopsis seedlings were grown at 22°C on solid growth media (0.5X MS basal salt medium with vitamins, 1% sucrose, pH 5.7 - 5.8 and 0.8% Agar (Sigma-Aldrich® A1296)) or liquid growth media (growth media without agar) and plants were grown in soil Sungro Sunshine® professional growing mix, under 16 hr-light/8 hr-dark cycles at 85-90 µmol $m^{-2}$ $s^{-1}$ light intensity. Previously described lines used in this work are: DR5::GUS [1], ARR7-OX (35S::ARR7) and ARR7 (D85N) [2], *tmkQ/tmk1; tmk2; tmk3; tmk4* [3–5], *tir1Qt*/*tir1afb1245/ tir1-1; afb1-3; afb2-3; afb4-8; afb5-5* mutant [6], *rop246/rop2; rop4; rop6* [7], ARR20-OX [8], DII-Venus [9], R2D2 [10], *shy2-3* [11], and $iaa18^{d}$/*iaa18-1* [12], *ech2-/-;ibr10-/-,* and *ech2-/-*; ECH2::YFP-ECH2 [13] and IBR10::YFP-IBR10 [13]. The *shy2-3* allele and $iaa18^{d}$ *-/+* (the homozygote is mostly sterile) are in the Ler-0 (Landsberg *erecta*) background. R2D2 reporter is in Col-utr (Columbia-Utrecht) background. All other mutants and reporters are in Col-0 ecotype (Columbia) background.

Seeds that were bleach-sterilized, vernalized for 2 days (see section on 'seed sterilization'), and grown under our growth conditions displayed distinctive phenotypic features at each timepoint that we analyzed: 0 HAP - swollen testa, 24 HAP - clear longitudinal rupture line in the testa; 36 HAP - protruded root tips but cotyledons remain inside the testa; 48 HAP - greening cotyledons emerging from the testa and root hair-like structures become noticeable at the root-hypocotyl junction; 60 HAP – further greening and oval-shaped opening cotyledons with a ~90° angle between them, and finally, 72 HAP – flat, green, open globular-shaped cotyledons with a ~180° angle between them.

#### Seed sterilization

Seed sterilization is crucial for the reproducibility of all experiments described in this manuscript. For routine experiments, sterilization was performed overnight by vapor-phase method (Clough and Bent, 1998) simply by placing seeds in a desiccator next to a beaker containing 80 mL of bleach and with 1 mL of concentrated HCl carefully added into the bleach. For experiments analyzing a specific developmental time point, seeds were bleach-sterilized as follows: 10 min agitation with 30% bleach in water with 0.1% TX-100 (Triton X-100), followed by rinsing 5 times with sterile water and vernalization for 48 h at 4°C in dark. Seeds were then plated on solid growth media and timing for Hours After Plating (HAP) or Days After Plating (DAP) begins.

**Chemical treatments**

All chemicals DMPNB-IAA [14], DMPNB-NAA [14], DMPNB-AcOH [14], Auxinole [15], BA (benzyladenine, CAS Number 1214-39-7), NAA (naphthalene-1-acetic acid, CAS Number 86-87-3, Sigma-Aldrich®), and IAA (indole-3-acetic acid, CAS Number 87-51-4, Sigma-Aldrich®) were added into warm media before pouring into plates or liquid media to obtain the final concentration indicated for each experiment from a 1000X stock solution in DMSO.

The 1000X stock solutions were prepared in small aliquots, carefully preserved in darkness at -20˚C, and discarded every month to avoid any potential effects of unintended uncaging/degradation due to light. Caged auxin and auxinole incubation treatments were performed in darkness with subtle motion.

**Microscopic analysis**

Imaging was performed with Zeiss LSM 880 (20X objective, NA= 0.8), Leica SP5 (20X objective, NA= 0.8), and/or Leica SP8 (40X objective, NA= 0.75) laser scanning confocal microscopes (LSCM). Confocal light path configuration was as follows: Venus, excitation 514 nm, emission 520-540 nm; ntdTomato, excitation 561 nm, emission 570-590 nm; propidium iodide (PI), excitation 514 nm, emission 540-600 nm; autofluorescence, excitation 355 nm, emission 400-600 nm; GFP, excitation 488nm, emission 500-530 nm.

**Imaging pavement cell outlines**

Pavement cell outlines were visualized by submerging seedlings for 10-30 seconds in 5-100 µg/mL propidium iodide (PI) (CAS Number 25535-16-4, Sigma-Aldrich®). For young cotyledon tissue (younger than 48 HAP), 5-25 μg/mL was used to avoid tissue damage and for green tissue (older than 48 HAP) 25-100 μg/mL was used. To avoid tissue damage, the delicate *tmkQ* and *tir1Qt III* cotyledons were stained as young cotyledons, regardless of their developmental age. Alternatively, cell outlines were obtained from cell autofluorescence under low UV laser excitation and 400-600 nm emission.

**Time-lapse tracking of auxin gradient in cotyledon**

The DII-Venus and mDII-Venus seeds were bleach-sterilized, plated on solid growth medium, and placed at 4°C for 48 hours of vernalization. After 18 hours in a growth chamber at 22˚C, the seed coat and the inner cotyledon (closest to the radicle) was removed with a syringe needle under the dissecting microscope, and the remaining cotyledon with hypocotyl and root was mounted on a coverslip with a block of solid medium for nutrients. Then, the coverslip was sealed in a moist box and imaged on the inverted LSCM ZEISS 880. The Venus images were obtained under the 512 nm 10% argon laser with the 512 nm Main Beam Splitter (MBS) Dichroic Mirrors and the 525 nm long pass filter for emission bandwidth. A full z-stack was performed for each cotyledon with 2-hour intervals under the same imaging settings. During the 2-hour imaging intervals, the seedlings were placed back in the growth chamber for cotyledon development. The z-stack images were subjected to the Maximum Intensity Projection processing for the final cotyledon Venus images. The DII-Venus or mDII-Venus intensities were measured within the circled ROIs in tip, middle, and basal regions of cotyledons Fiji [16] (Fiji Is Just ImageJ).

**Computational analysis of pavement cell shape**

Morphometric analysis of pavement cells was performed with PaCeQuant [17] in batch mode, following default settings and all recommendations suggested in published literature [18]. The initial data quality analysis is performed with PaCeQuantAna, a very useful Rstudio-compatible library package for PaCeQuant data analysis [19]. However, the final figures result from custom R code implemented to concatenate the result tables from each image into a data frame for subsequent graphing in violin, split-violin, or scatter plots using ggplot2. This code can be found at https://github.com/pperezh/global-local-auxin.

Margin Roughness heatmaps of entire cotyledons were performed with PaCeQuant in 3 steps. First, images are processed in SEGMENTATION_ONLY mode. If segmentation crashed, we use inverted gray LUT, adjusted contrast to highlight borders, and/or re-adjusted the segmentation filter configuration to WATERSHED with a NIBLAK threshold value of 1 (the preset value is 4). The resulting ROI list is edited to remove unwanted cells such as marginal cells, guard mother cells, or guard cells. Second, the edited ROI list was processed again with PaCeQuant on FEATUES_ONLY mode. Third, the resulting feature data was uploaded to the "FeatureColorMapper" add-on tool for PaCeQuant [19], selecting Margin Roughness as the feature to be mapped and using colors blue (#0000FF) for minimum and orange (#FF8C0) for maximum values, without any cell size filter.

**Single cell tracking of *de novo* lobe formation**

For *de novo* lobe formation, seeds were bleach-sterilized and vernalized for 2 days. Seedlings were grown for 3 days on solid growth media, carefully submerged in propidium iodide (10-30 sec in 100 µg/mL PI), and intact seedlings were mounted in water exposing the adaxial epidermis for imaging with confocal microscopy (pre-treatment image). Then, seedlings were transferred to liquid media containing mock or auxin treatment for the indicated time (2.5 days) with constant subtle motion in light. After treatment, seedlings are carefully manipulated for re-imaging the adaxial epidermis (post-treatment image). Single cell tracking was performed by manually matching images pre-and post-treatment. Images capturing multiple cells (350 µm x 350 µm) were preferred to increase the points of reference for manual matching. Segmentation was performed manually for posterior analysis with PaCeQuant.

**Auxin uncaging**

Careful handling of seedlings is extremely important for uncaging experiments. All uncaging experiments (except on 24 HAP R2D2 cotyledons) were carried out on intact seedlings (without cotyledon dissection), therefore, the adaxial epidermis needed to be accessible. This is only possible in seedlings after 72 HAP. Uncaging experiments required the following steps: 1) seedlings were incubated for 5 h in liquid media supplemented with DMPNB-IAA, DMPNB-NAA or DMPNB-AcOH in subtle motion and darkness; 2) suitable healthy looking seedlings were carefully mounted to capture a pre-UV image using propidium iodide or autofluorescence; 3) a UV pulse treatment was applied to a region of interest (355 nm, 30 s, 60 mW at 1% or 25% laser); 4) if applicable, seedlings were carefully placed back on solid media for a post-UV incubation period; and 5) a post-UV single or time-lapse image was acquired.

Uncaging occurs when UV light breaks the bond between DMPNB and the caged molecule [14] (Figure S2C). Caged auxin solutions are stable under -20˚C in the dark. Light from normal overhead fluorescent lamps (10-20 W) do not decompose caged molecules, but direct sunlight exposure will. The UV light treatment is restricted to defined regions of interest (ROIs) within the cotyledons by digital zoom on the microscope. ROIs can contain single cells (20 µm x 20 µm) or multiple cells (85 µm x 85 µm). As a reference, the size of the multiple cell ROI used for all experiments with 24 HAP R2D2 cotyledons was 85 µm x 85 µm. The precision of the UV-light treated areas was evaluated by simply examining autofluorescence, as there is a clear signal bleaching after UV exposure (Figure S2G-top panel). UV light treatment was lethal above 25% laser power, determined by propidium iodide as a cell viability marker. Although the laser 355 nm was set to 60 mW for all experiments, the laser percentage used varies according to the experiment.

*24 HAP cotyledons*

Caged auxin in the nanomolar range was used for assays with matured embryos (24 hours after plating) before cuticular formation. Older cotyledons render very weak basal DII-Venus because endogenous auxin levels dramatically increase after 24 HAP. These assays aim to detect protein degradation within a few minutes. In experiments with the more fragile R2D2 dissected cotyledons at 24 HAP, the laser power was set at 1%. With higher laser intensities, strong signal interference was captured (Figure S2G-bottom panel), where 1% and 5% laser intensities are compared. 24 HAP RDD2 seedlings were incubated with 50 nM DMPNB-IAA for 5h in darkness, carefully dissected, UV-treated, and imaged every 15 sec for

5 min after uncaging. Time-lapse imaging starts after 30 sec of UV irradiation, which is the time necessary to set up the time-lapse.

*3.5 DAP cotyledons*

Caged auxin concentration in the micromolar range was used for assays with older seedlings (3.5 DAP) as the epidermal cuticle has formed making older cotyledons less permeable to caged auxin. In experiments involving "open cotyledons" meaning 72 HAP (3 DAP) or older, we used 25% laser power. Non-dissected cotyledons underwent either mock treatments with 100 μM of DMPNB-AcOH (acetic acid) or caged auxin 100 μM DMPNB-NAA for 5 h in darkness. The cotyledons were then imaged pre-UV, UV-treated, and incubated in solid media for the indicated time (15h or 2 days in darkness) after UV treatment. Finally, cotyledons were dissected and imaged by confocal microscopy.

**GUS (beta-glucuronidase) activity assays**

Histological assays (GUS staining with X-gluc as substrate) were performed with slight modifications from previous reports [20,21]. In brief, plant samples were fixed in cold 80% Acetone for 2 h, then vacuum infiltrated with GUS solution for 10 min and incubated in the dark at 37˚C for 0.5h. Alternatively 2-4 h incubation without vacuum infiltration. For the cytokinin treatment experiments, the incubation time was doubled for a clearer visualization of the cytokinin suppressive effect. After staining, plant tissues were cleared in 70% ethanol until chlorophyll was not detected.

Fluorometric assays, (GUS activity in intact plant tissue with 4-MUG as substrate) were performed with adaptations from previous reports [22,23]. Samples were incubated in 96-well microplates (one sample per well) with 150 μL lysis buffer (50 mM sodium phosphate, pH 7.0, 10 mM EDTA, 0.1% Triton X-100). For this assay, we define a sample as a pair of cotyledons from a seedling. The sample must be completely submerged in the buffer for substrate penetration. The reaction is then started with 1 mM 4-MUG, incubated at 37˚C for 90 min, and then stopped with 50 μL of 0.2 M $Na_2CO_3$. The fluorescence intensity of the reaction product, 4-MU, was directly measured with the multi-mode plate reader SpectraMax iD5 using the fluorescence intensity read mode under the following specific wavelength; 365 nm for excitation and 455 nm for emission. A standard curve elaborated with 4-MU (in lysis buffer plus stop solution) corroborated linearity R2 = 0.99 for this assay in our conditions between 10 nM and 1000 nM of 4-MU. GUS activity is expressed as molar units of 4-MU nm per sample.

**ROP2 and ROP6 activity assays**

ROP2/6 activity measurements were performed as described previously [3,5]. For this, 35S::GFP-ROP2 and 35S::GFP-ROP6 seedlings were grown for 4 days in a 6-well cell culture plate (BEAVER™) in liquid growth medium (as defined in growth conditions) and later grown for 2 additional days in the presence of 200 μM auxinole to completely inhibit TIR1-AFB signaling. Then the cultured seedlings (about 20 seedlings per well) were treated in 200 μM auxinole with or without 1 μM NAA for 5 minutes, dried, and frozen immediately in liquid nitrogen. A total of 0.2-0.5 g seedlings were collected for western blot-based ROP activity measurements. Active ROP proteins (GTP-bound ROP) were pulled down by 100 μL MBP-RIC1 beads (3h at 4°C) and detected by western blot analysis using anti-GFP (Santa Cruz Biotechnology) and Goat Anti-Mouse IgG-HRP (Bio-Rad). Total ROP was determined before MBP-RIC1 incubation. ImageJ was used to quantify ROP activity, defined as the ratio between active and total ROP.

**qRT-PCR analysis**

For the detection of basal expression levels, 30-40 cotyledons were quickly dissected from 4.5-day-old seedlings. For detection of expression levels in response to auxin, 30-40 4.5-day-old seedlings were treated for 0.5h in a liquid medium with the indicated concentration of NAA. Only the *tir1Qt III* seedlings were collected for both experiments. After collection, the tissue was quickly dried and immediately frozen in liquid nitrogen. Then, samples were ground using TissueLyzer® (Qiagen), and total RNA was isolated with Rneasy Plant Mini Kit (Qiagen) using the in-column DNAse I treatment (Qiagen). Reverse transcription was performed using iScript cDNA synthesis kit (Bio-Rad). Primers were designed with Primer3 spanning an intron whenever possible. Primer pairs used for the target genes were: ECH2-F (5’-AACCAGGGCTTAGCTGTGAA) and ECH2-R (5’-GGTTTTGACAGCAGTTGGGT), IBR3-F (5’-

GACATGCGGACTCCAGGTAT) and IBR3-R (5'-TTAGCTCCATTCCACGCTCT), IBR10-F (5'-TCGATGCCGGATTATTTCTC) and IBR10-R (5'-ATCGACGCATAAACCTCACC). Primer pairs used for reference genes were: UBQ5-F (5'- AGCCGAAGAAGATCAAGCACA) and UBQ5-R (5'-GCCATGAAAGTCCCAGCTCC), AP4M-F (5'-GCTGGACGTCTTAAGGCTGA) and AP4M-R (5'-GCAGCACCGGGTTCTAACTC), PP2AA3-F (5'-TGTTCCAAACTCTTACCTGCGG) and PP2AA3-R (5'-ATGGCCGTATCATGTTCTCCAC). Each primer pair was analyzed by RT-PCR and melting curve analysis detecting the amplification of one single amplicon. qRT-PCRs were conducted with iQ SYBR Green Supermix (Bio-Rad) using the LightCyler98 (Roche) or the CFX384 (Bio-Rad) for fluorescence reading. Primer efficiency was calculated based on the raw fluorescence data using linear regression on each reaction with the software LinReg [24]. Five reference genes: *UBQ5* (AT3G62250), *PP2AA3* (AT1G13320), *AP4M* (AT4G24550), *EF1A* (AT5G60390), *UBC9* (AT4G27960) were initially analyzed with geNorm [25]. Although all genes showed acceptable geNorm M values, we selected the 3 most stable ones in our sample group: *UBQ5*, *PP2AA3*, and *AP4M*. Expression levels are normalized to the geometric mean of the calibrators and relativized to values obtained in mock conditions. Relative expression is processed with the software qBase+ [26]. Three biological replicates were used, and each sample was run with 3 technical replicates.

**Genotyping**

The mutant *tir1Qt*/*tir1-1;afb1-3;afb2-3;afb4-8;afb5-5* was genotyped using the following primers: *afb1-3_LP*, AACGGAAGACTAGGAAGCGAG, *afb1-3_RP* (5'-GCAACAGCTTCAAGACCTTTG, *afb2-3_LP* (5'-TCAACGGTCAAGATCCATCTC), *afb2-3_RP* (5'-CTGCAATTAGCGGCAATAGAG), *afb3-4_LP* (5'-TCATGTTGCTTACAAATTGCG), *afb3-4_RP* (5'-TCTGCAAACAGATGACAAACG), *afb4-8_LP* (5'-CTAAACACAGTAAGCTGCGGG), *afb4-8_RP* (5'-CCTAATTGGGGAGCTCAATTC), *afb5-5_LP* (5'-GTTGGATCTACCCTCTACCGC), *afb5-5_RP* (5'-GTGGCAATTGAGTATGATGGG). The mutation in *tir1-1* was identified by Sanger sequencing as a point mutation (GGT > GAT) resulting in a G→D change at amino acid 147, as described previously [6]. The mutant *tmkQ/tmk1;tmk2;tmk3;tmk4* was genotyped for *tmk1-/-* (SALK_016360), *tmk2 -/-* (SAIL_1242_H07), *tmk3-/-* (SALK_129759), and *tmk4-/-* (GABI_191D02) using PCR, as previously described [4]. Mutation in *iaa18*$^{d}$*/iaa18-1* was identified by sanger sequencing of a *iaa18d-F2*/*iaa18d-R* PCR product or by digestion of the *iaa18d-F1*/*iaa18d-R* PCR amplicon with the restriction enzyme FokI, as described previously [12]. Primer sequences as follows: *iaa18d-F1* (5'-ATCTCTCCGAAGCTGCTTGAC), *iaa18d-F2* (5'-TACTGCCCTAGTTTTCCTCTG), *iaa18d-R* (5'-CATGCCTCCTTGTCTCTTTGG).

**Transgenic rescue of mutants**

*tmkQ* rescue

The pGWB504/TMK1::TMK1-sGFP plant expression vector was generated as follows: The *TMK1* sequence, including the 3.5 kb region including the presumptive promoter and genomic coding region without the stop codon, was amplified using the primers *TMK1-F* (5' GGGGACAAGTTTGTACAAAAAAGCAGGCTCCCGTTTGGAACAGATCTTTCAGCTATACAT) and *TMK1-R* (5' GGGGACCACTTTGTACAAGAAAGCTGGGTCTCGTCCATCTACTGAAGTGAATGACTCTGC), and then inserted into the pDONR207 vector. The resulting entry vector was then transferred into the Gateway®-compatible binary vector pGWB504 with a C-terminal sGFP tag via Gateway® recombination with the enzyme LR Clonase® II Plus (Invitrogen). The recombinant vector pGWB504/ TMK1::TMK1-sGFP was sequence-verified and subsequently transformed into *tmk1-/-;tmk2-/-;tmk3-/-;tmk4-/+* plants using the standard Agrobacterium tumefaciens-mediated floral dip method. T1 seeds were screened with 30 µg/mL Hygromycin B to confirm the presence of the pGWB504/ TMK1::TMK1-sGFP construct. Then, *tmkQ* rescue lines (*tmkQ*; TMK1::TMK1-GFP) were obtained by confirming *tmk* mutations in the hygromycin-resistant T1 plants for *tmk1-/-* (SALK_016360), *tmk2-/-* (SAIL_1242_H07), *tmk3-/-* (SALK_129759), and *tmk4-/-* (GABI_191D02) using PCR. Finally, T2 seeds were use for the PC analysis.

*ech2-/-;ibr10-/-* rescue

*ech2-/-;*ECH2::YFP-ECH2 and IBR10::YFP-IBR10 plants were crossed to *ech2-/-;ibr10-/-*. F2 seedlings were screened by fluorescent signal in roots. YFP+ F2 plants were then genotyped for *ech2-/-* and *ibr10-/-* by PCR and/or enzymatic digestion. PC assays and protein expression analysis were performed on F3 seeds.

**Graphs and statistical analysis**

All graphs were constructed using Microsoft Excel®, GraphPad® Prism 9, or ggplot2 in RStudio. Statistical analysis was performed in ggplot2 or GraphPad® Prism 9 using Student's *t*-test (parametric) and Wilcox test (non-parametric).

**Image Processing**

Images were processed with proprietary software Zeiss ZEN® 3.7 software, Leica Application Suite® X software, or the freeware Fiji (Fiji is Just) ImageJ v1.54. Most pavement cells images are displayed as inverted gray look-up table (LUT). Final multi-panel figures were built using the freeware Inkscape v1.3.

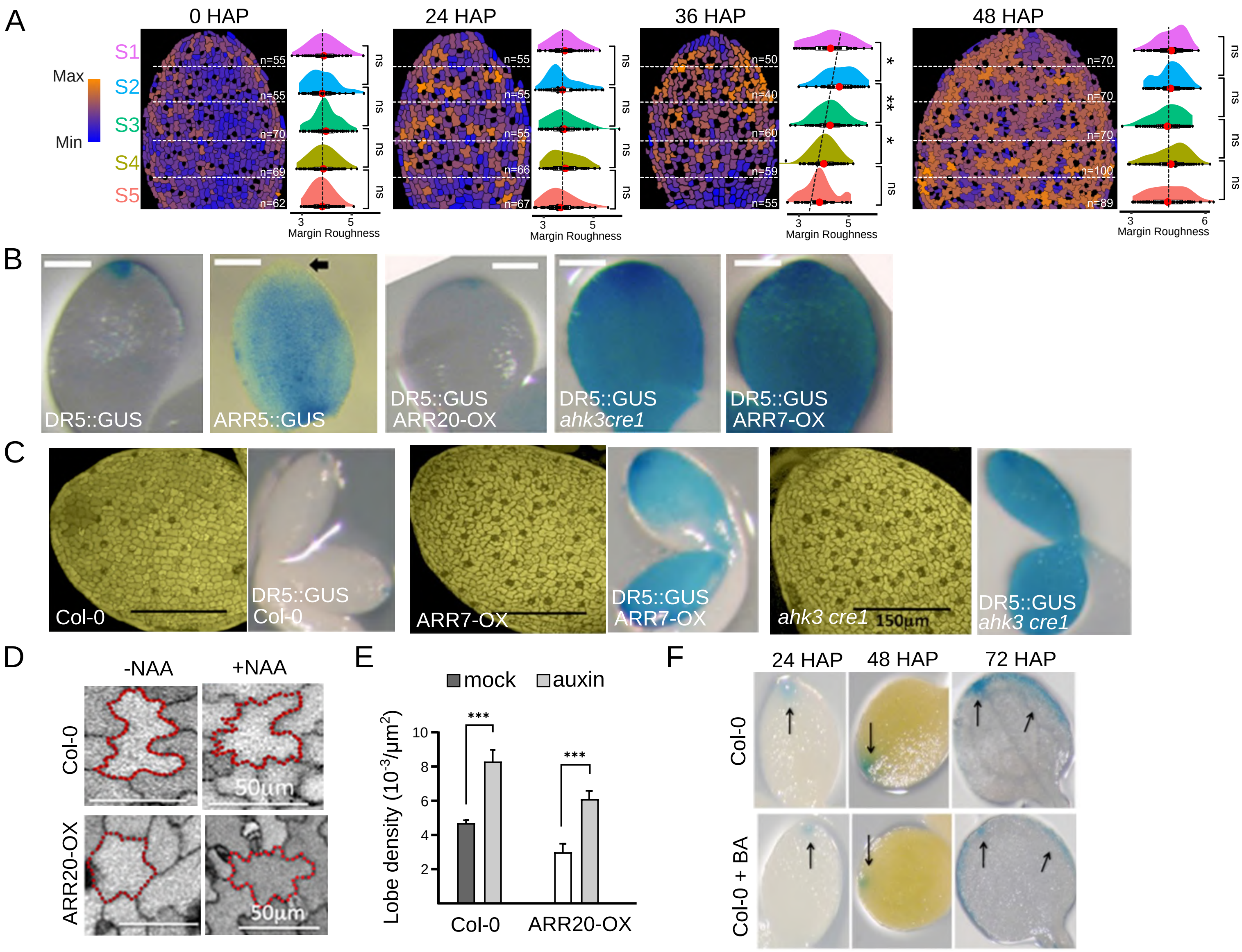
A
0 HAP
24 HAP
36 HAP
48 HAP
Max
Min
S1
S2
S3
S4
S5
n=55
n=55
n=70
n=69
n=62
n=55
n=55
n=55
n=66
n=67
n=50
n=40
n=60
n=59
n=55
n=70
n=70
n=70
n=100
n=89
ns
*
**
Margin Roughness
B
DR5::GUS
ARR5::GUS
DR5::GUS ARR20-OX
DR5::GUS ahk3cre1
DR5::GUS ARR7-OX
C
Col-0
DR5::GUS Col-0
ARR7-OX
DR5::GUS ARR7-OX
ahk3 cre1
150µm
DR5::GUS ahk3 cre1
D
-NAA
+NAA
Col-0
ARR20-OX
50µm
E
mock
auxin
Lobe density (10^-3/µm^2)
***
Col-0
ARR20-OX
F
24 HAP
48 HAP
72 HAP
Col-0
Col-0 + BA

**Figure S1. Cytokinin regulates the spatiotemporal pattern of PC interdigitation in cotyledons by acting as a developmental break and suppressing auxin transcriptional response.**
(A) Heat map of Margin Roughness (MR) values for pavement cells within each indicated segment of the entire cotyledons at different HAP. Cotyledons were dissected before imaging: swollen seed (0 HAP), broken testa (24 HAP), emerged radicle (36 HAP), and green open cotyledon (72 HAP). n=40-100 cells per segment as indicated on each segment. Dashed vertical lines indicate MR trend along the proximo-distal axis. *t*-test, ns=no significance, *$p<0.05$, **$p<0.01$.
(B) Cytokinin antagonizes auxin in early developing cotyledons. The cytokinin signaling marker *ARR5::GUS* shows a complementary spatial expression pattern with *DR5::GUS*. Additionally, *DR5::GUS* expression was reduced in cotyledons over-expressing *ARR20* (*ARR20-OX*), a positive regulator of cytokinin signaling, but was expanded to the entire cotyledon in cotyledons over-expressing *ARR7* (*ARR7-OX*), a negative regulator of cytokinin signaling or in the loss-of-function *ahk3cre1* mutants for cytokinin perception. Cotyledons were analyzed at 24 HAP. Scale bars = 50 µm.
(C) Inactivation of cytokinin signaling caused constitutive activation of PC interdigitation in un-germinated cotyledons. In *ARR7-OX* and *ahk3 cre1* lines, PCs became constitutively interdigitated throughout the whole cotyledons prior to seed germination, consistent with *DR5::GUS* expression. Scale bars =150 µm.
(D-E) NAA restored PC interdigitation defects observed in *ARR20-OX*. (D) Representative images of pavement cells from seedlings treated with mock or exogenous 20 nM auxin NAA both in wild-type *Col-0* and *ARR20-OX* cotyledons. PC shapes were analyzed by autofluorescence using a 400-600 nm bandwidth in confocal microscopy at 48 HAP. Bars = 50 µm. (F) Quantitative analysis of images in D. Numbers of lobes were manually counted and normalized per 2D cell area. Lobe density doubled after NAA treatments in both wild-type *Col-0* and *ARR20-OX* cotyledon. *t*-test *** $p<0.001$.
(F) Cytokinin reduces the *DR5::GUS* signal at the tip and margin of cotyledons in germinating seedlings. Treatment with exogenously supplemented cytokinin 50 nM BA (benzyladenine) reduced GUS staining at various stages of cotyledons. Bars =50µm.

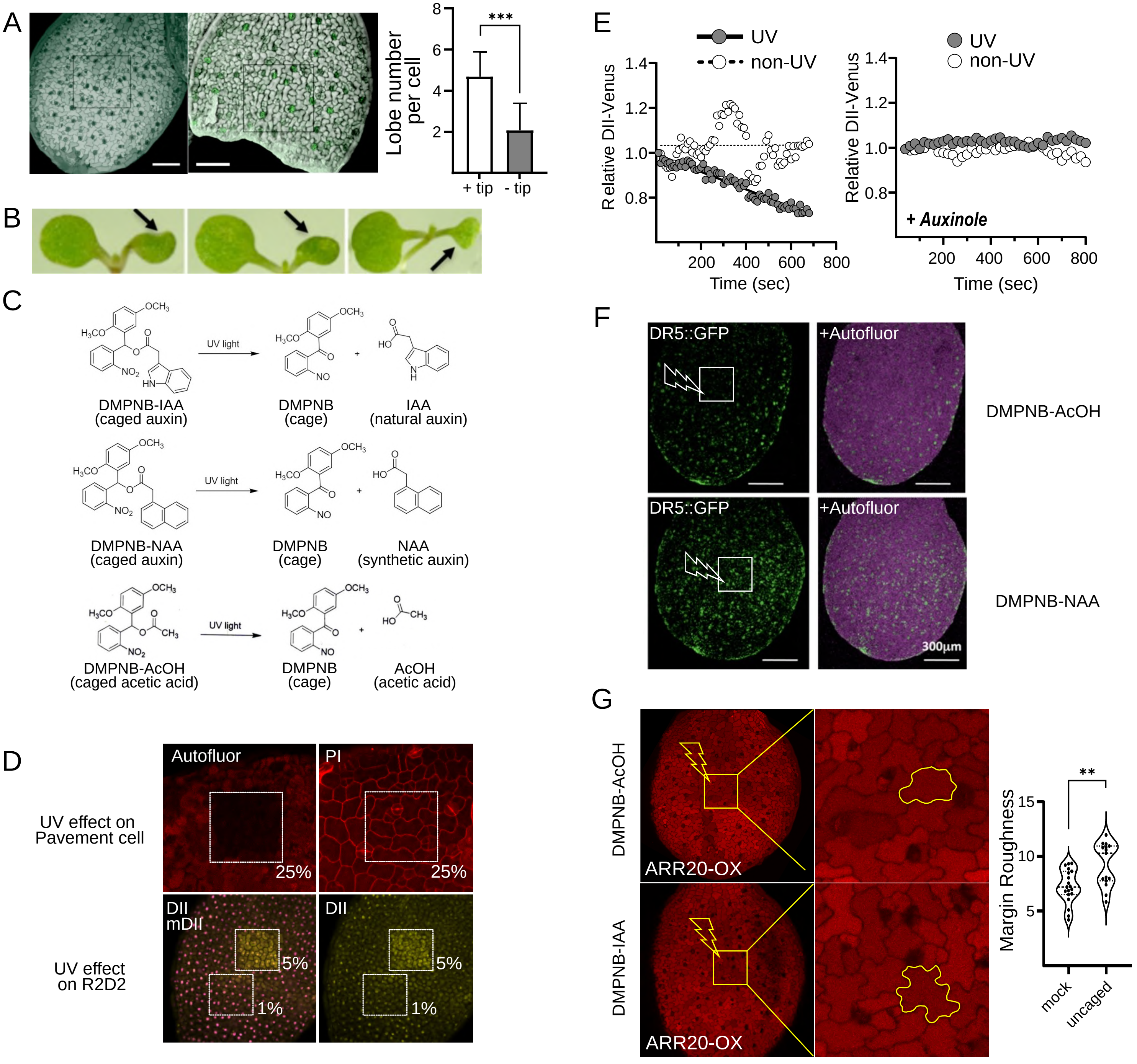
A
Lobe number per cell
***
8
6
4
2
+ tip
- tip
B
C
OCH3
H3CO
NO2
HN
UV light
DMPNB-IAA
(caged auxin)
DMPNB
(cage)
IAA
(natural auxin)
DMPNB-NAA
(caged auxin)
NAA
(synthetic auxin)
DMPNB-AcOH
(caged acetic acid)
AcOH
(acetic acid)
D
UV effect on
Pavement cell
Autofluor
PI
25%
UV effect
on R2D2
DII
mDII
5%
1%
E
UV
non-UV
Relative DII-Venus
1.4
1.2
1.0
0.8
200
400
600
800
Time (sec)
+ Auxinole
F
DR5::GFP
+Autofluor
DMPNB-AcOH
DMPNB-NAA
300μm
G
DMPNB-AcOH
DMPNB-IAA
ARR20-OX
Margin Roughness
**
15
10
5
mock
uncaged

**Figure S2. Assessing the presence of a global auxin signal by surgical removing cotyledon tips and auxin uncaging**

(A) Removal of the cotyledon's tip inhibited PC interdigitation. (Left) Representative images of cotyledons displaying PC shape at 48 hours after plating (48 HAP). mTalin-GFP in cotyledons without removing the tip (+tip) and after removing the tip (-tip) at 24 HAP, showed that cotyledon cells were alive and expressing GFP after removing the tip. (Right) Quantification of lobes per cell on intact cotyledon (+tip) and severed cotyledon (-tip). n > 200 cells, from 10 seedlings. *t*-test, *** p-values<0.001.

(B) Cotyledons from 96 HAP seedlings after tips or margins were surgically removed at 24 HAP.

(C) Chemical reactions for uncaging auxin or its analogs. After UV treatment, caged IAA (DMPNB-IAA) releases IAA from the caging group DMPNB (2,5-dimethoxyphenyl)(2-nitrobenzyl), whereas caged NAA (DMPNB-NAA) released NAA and DMPNB. Caged acetic acid (DMPNB-AcOH) is used as a mock treatment to normalize the effects of photolyzed by-products from the DMPNB caging group on physiological events. Stock solutions 10-50 mM in DMSO were used. Stock solutions are stable at 20°C in the dark. Light from normal fluorescent lamps (10-20 W) does not decompose caged molecules, but direct sunlight exposure will.

(D) Top panel - UV effect on Pavement cells. The accuracy of the UV irradiated region was corroborated by the bleached signal autofluorescence pattern (Autofluor). The effect of the UV irradiation on cell viability was evaluated by vital marker propidium iodide (PI). Cell viability remained intact with intensity below 25% of the 355nm laser at 60 mW. Bottom panel - UV effect on R2D2. The DII-Venus channel, and not the mDII-Venus channel displays an interference signal with the 355 nm laser at intensities higher than 1%. Therefore, uncaging experiments in R2D2 cotyledons of 24 HAP embryos were performed at 1%, although cell viability remains intact with higher laser intensities <25%.

(E) Effective release of caged auxin. Auxin uncaging induced a decrease in the DII-Venus signal at a single-cell level, whereas auxinole abolished this uncaging-induced change. Venus nuclear signal intensity was quantified over 13 minutes after uncaging in both UV-treated (gray) and non-UV-treated cells (white). In this experiment, R2D2 seeds were grown for 24 HAP, seed coats were removed, and seedlings were incubated with 20 μM DMPNB-IAA in the absence or presence of auxinole, a potent TIR1/AFBs-Aux/IAA interaction inhibitor. The graph showes the average value from 5 different uncaging events. The black line is the linear model for UV-treated cells. Similar results in 2 experimental replicates.

(F) Local auxin uncaging induced *DR5::GFP* levels in the whole cotyledon. DR5::GFP seeds were germinated in 10 μM BA Cytokinin for 2 days to suppress DR5::GFP expression, and then seedlings were bathed in liquid media containing 100 nM caged NAA or caged control in darkness for 5 hours. Caged compounds were released by UV irradiation (indicated by squares) and images were taken 20 hours after uncaging.

(G) Uncaged auxin-induced PC interdigitation in the uncaging region of cotyledons. ARR20-overexpressing seedlings (3.5 DAP) were incubated either in 20 μM of DMPNB-AcOH (acetic acid/mock) or 20 μM DMPNB-IAA (caged auxin) for 5 h in darkness, then UV-irradiated for 30 sec (square box, 25% laser, 60mW). (Left) 15 hours after UV irradiation, cotyledons were dissected and imaged with confocal microscopy using the autofluorescence spectrum (emission 400-600nm). (Right) Margin Roughness of PCs within the UV-irradiated region was analyzed using PaCeQuant after manual segmentation, n= 15-19 cells, analyzed from 5 cotyledons per treatment. Similar results were obtained in 3 experimental replicates. Similar results were obtained with 2 μM DMPNB-IAA. Scale bar = 100 μm. *t*-test, ** p-values<0.01.

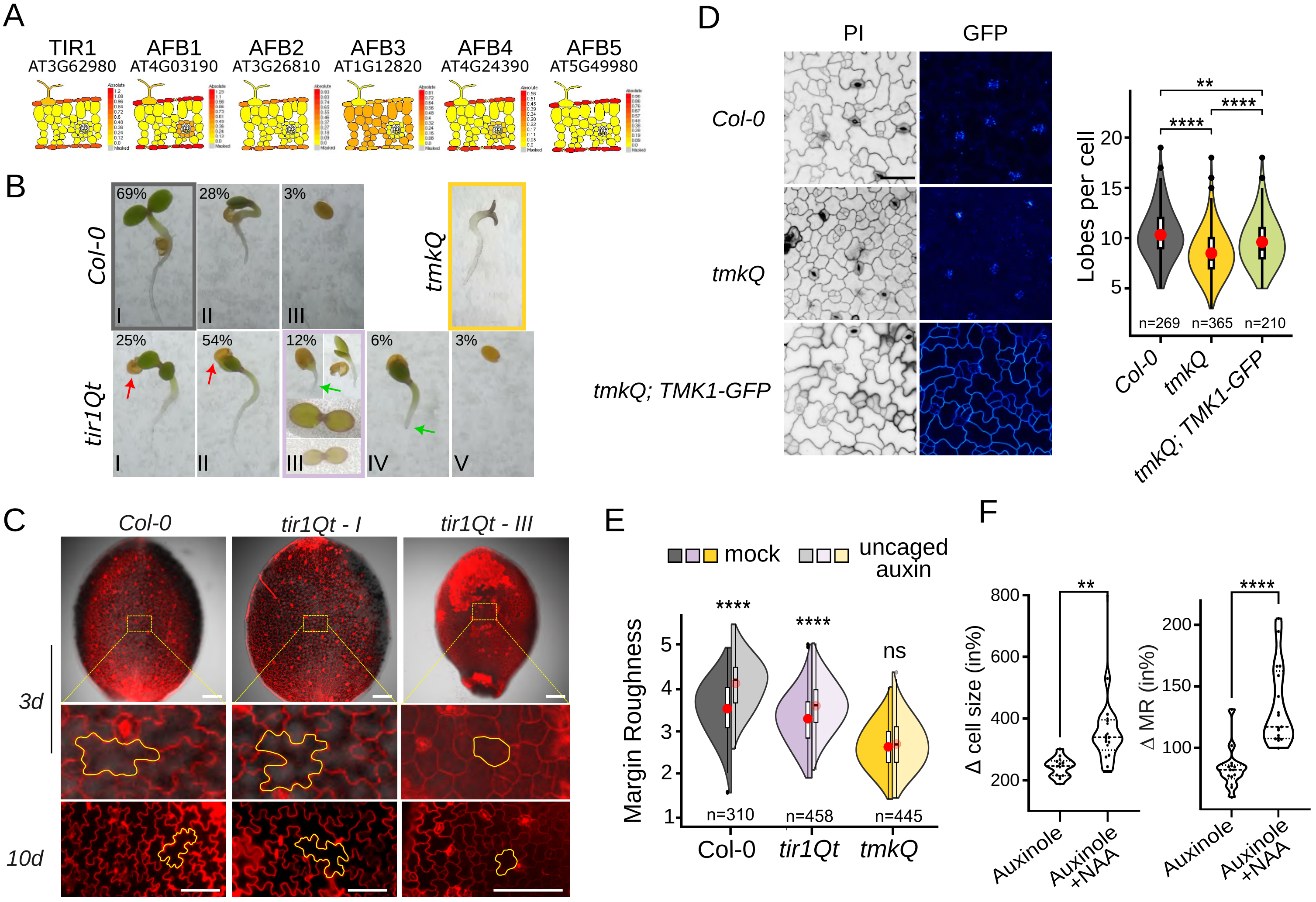
A
TIR1
AT3G62980
AFB1
AT4G03190
AFB2
AT3G26810
AFB3
AT1G12820
AFB4
AT4G24390
AFB5
AT5G49980
B
Col-0
69%
28%
3%
I
II
III
tmkQ
tir1Qt
25%
54%
12%
6%
3%
I
II
III
IV
V
C
Col-0
tir1Qt - I
tir1Qt - III
3d
10d
D
PI
GFP
Col-0
tmkQ
tmkQ; TMK1-GFP
Lobes per cell
**
****
****
n=269
n=365
n=210
Col-0
tmkQ
tmkQ; TMK1-GFP
E
mock
uncaged auxin
Margin Roughness
****
****
ns
n=310
n=458
n=445
Col-0
tir1Qt
tmkQ
F
Δ cell size (in%)
**
Auxinole
Auxinole +NAA
Δ MR (in%)
****
Auxinole
Auxinole +NAA

**Figure S3. TIR1/AFBs and TMK auxin signaling pathways function in PC interdigitation.**

(A) Gene expression from cell-type specific translatome data obtained at http://efp.ucr.edu for TIR1 (At3g62980), AFB1 (At4g03190), AFB2 (At3g26810), AFB3 (At1g12820), AFB4 (At4g24390), AFB5 (At5g49980). Graphs are obtained from datasets called "relative" because these values are best to evaluate cell type-specific differences [27].

(B) Phenotype of 3 days-old seedlings of wild type Col-0, and loss-of-function mutants *tir1Qt (tir1afb1245)* and *tmkQ.* The phenotypes of *tir1Qt* siblings are notoriously widely varied. For example, *tir1Qt* class I and II siblings are unable to detach the seed testa (red arrow), whereas *tir1Qt* class III and *tir1Qt* IV siblings are unable to establish a root (green arrows). Auxin responsive analysis in Figure 3 and Figure S3E corresponds to seedlings wild type Col-0 (gray), *tir1Qt* class III (purple) and *tmkQ* (yellow). The occurrence frequency for each phenotype is annotated in the upper left as a percentage, n=208 seedlings.

(C) PC phenotype of wild type *Col-0* and *tir1Qt* classes I and III at 3 DAP (3d, upper panel) and 10 DAP (10d, lower panel). PC shapes (yellow) in the middle panel show the strong PC defects of *tir1Qt* III. At 10d, *tir1Qt* I displayed big flat cotyledons and big interdigitated PCs, however, *tir1Qt* III exhibited strong PC defects.

(D) TMK1-GFP rescued *tmkQ* PC phenotype. Representative images of PC shape and green fluorescence (GFP) from 3-day-old seedlings of wild type Col-0, quadruple mutant *tmkQ,* and the rescue line *tmkQ; TMK1-GFP*. The rescue line was obtained by transformation of TMK1::TMK1-GFP into the quadruple mutant *tmk1-/-;tmk2-/-;tmk3-/-;tmk4-/+*. T2 seeds were screened with Hygromycin B. Resistant plants were screened by PCR for *tmk-/-* mutants. Images are obtained with genotyped, stable, T3 seedlings TMK1-GFP/*tmkQ.* The box plot inside each violin plot depicts four quartiles and the median. The red dot depicts the average. n=210-365 cells analyzed from 8 different cotyledons. Results are representative of 3 experimental replicates. Scale bar= 50 µm *t*-test, **p<0.01, ****p<0.0001.

(E) PC interdigitation analyzed by Margin Roughness from the auxin responsiveness experiment shown in Figure 3A. Seedlings were grown for 5 days on liquid medium in mock (DMSO 0.01%) or 20nM NAA treatment. Col-0 (gray), *tir1Qt III (purple),* and *tmkQ* (yellow). Split violins show for each genotype the mock (opaque) and NAA treatment (translucent) values. The box plot inside each violin plot depicts four quartiles and the median. The red dot depicts the average. n is indicated below each violin plot, coming from at least 8 different cotyledons, each from different seedlings. Similar results were obtained in 5 independent experiments. *t*-test, ns=no significant, ****p<0.0001.

(F) Percentage variation of pavement cell size (%Δ cell size) and Margin Roughness (%ΔMR) from single-cell tracking experiment shown in Figure 3C. Percentage variation in the cell size and margin roughness (MR) was calculated with manually paired images captured before (3 DAP) and after treatment (+2.5 days of treatment). n=28-34 cells, *t*-test, *p<0.05, **p<0.01, ***p<0.001, ****p<0.0001.

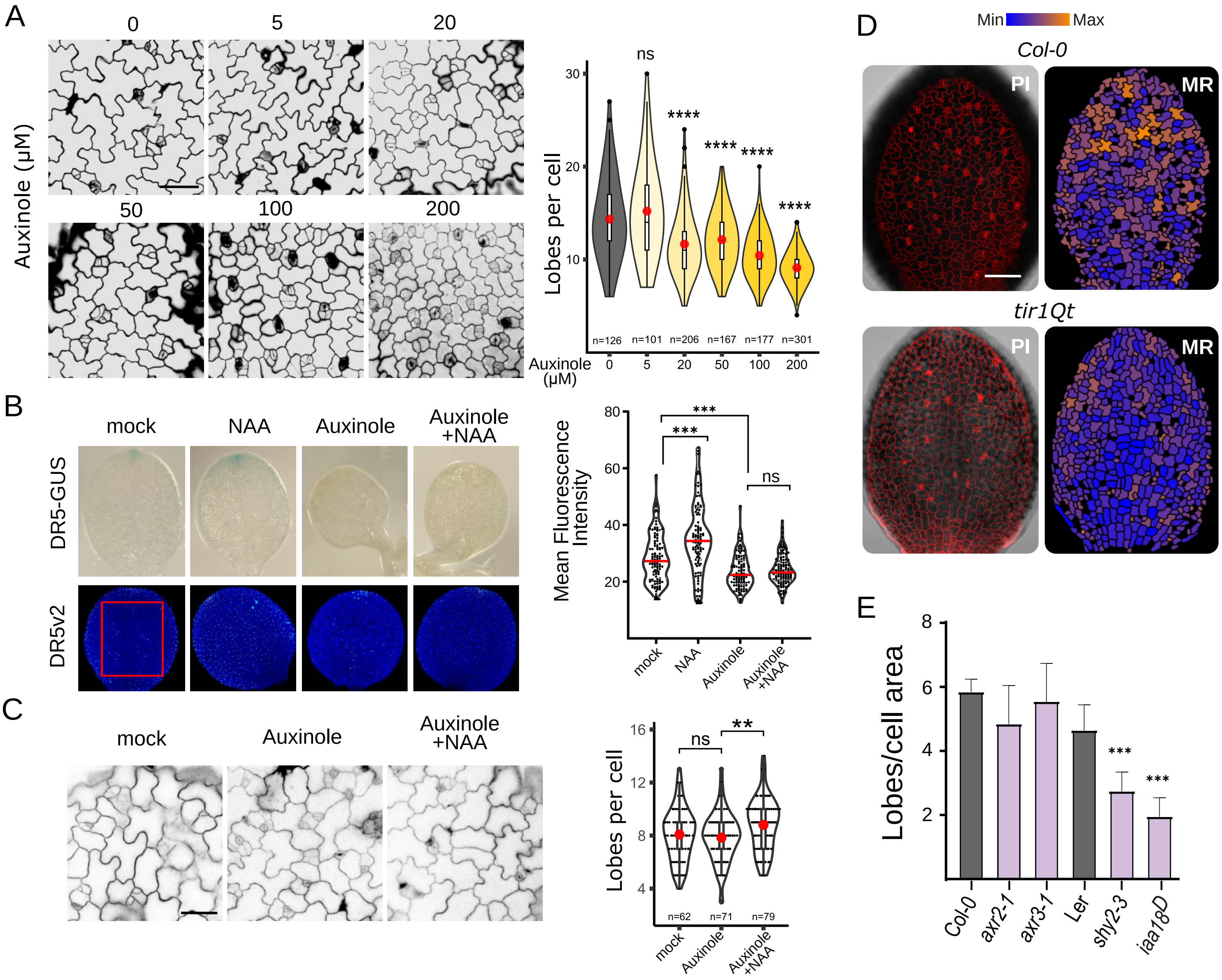
A
Auxinole (µM)
0
5
20
50
100
200
Lobes per cell
ns
****
****
****
****
n=126
n=101
n=206
n=167
n=177
n=301
Auxinole (µM)
0
5
20
50
100
200
B
mock
NAA
Auxinole
Auxinole +NAA
DR5-GUS
DR5v2
Mean Fluorescence Intensity
***
***
ns
mock
NAA
Auxinole
Auxinole +NAA
C
mock
Auxinole
Auxinole +NAA
Lobes per cell
ns
**
n=62
n=71
n=79
mock
Auxinole
Auxinole +NAA
D
Min
Max
Col-0
PI
MR
tir1Qt
PI
MR
E
Lobes/cell area
***
***
Col-0
axr2-1
axr3-1
Ler
shy2-3
iaa18D

**Figure S4. TIR1/AFB-based auxin signaling regulates PC interdigitation.**

(A) Dose-dependent effect of auxinole on PC interdigitation. Wild-type Col-0 seeds were germinated in the absence of auxinole for 6h, and germinated seedlings were grown for 4 days in a solid medium supplemented with increasing concentrations of auxinole from 5 to 200 µM. Images are obtained with confocal microscopy after staining cotyledons with 50 µg/mL propidium iodide. Lobes per cell were obtained with the software PaCeQuant. The box plot inside each violin plot depicts four quartiles and the median. The red dot depicts the average. n =101-301 cells as indicated below each graph from 10-12 cotyledons, each from a different seedling. Similar results were obtained in 3 independent experiments. Scale bar =50 µm. *t*-test, ns=no significant, ****p<0.0001.

(B) auxinole inhibited transcriptional response induced by exogenous auxin. (Left) Representative images from 3-day-old *DR5::GUS* and *DR5v2* seedlings that were treated in darkness for 72h and 24h, respectively, in liquid medium containing mock (DMSO 0.01%), 100 nM NAA, 200 µM auxinole, or 200 µM auxinole + 100 nM NAA (with a 30min auxinole pre-treatment). (Right) Quantification of nuclear DR5v2 signal intensity in the center of epidermis field (red ROI). The violin plot depicts median values in red. n >102 cells from 5-7 cotyledons, each from a different seedling. Similar results were obtained in 4 independent experiments. *t*-test, ns=no significant, ***p<0.001.

(C) Auxin-induced PC lobing in *tir1Qt* in the presence of auxinole. To ensure that any residual TIR1/AFB signaling activity in *tir1Qt III* was completely eliminated, *tir1Qt III* seedlings were grown for 3 days in a liquid medium and then treated with 100 nM NAA for 48 h in the presence or absence of 200 µM auxinole. Co-treatments were preceded by a pre-incubation with auxinole for 30 min. Then, cotyledons were stained and imaged with PI and quantified with PaceQuant. The red dot depicts average values. The number of cells analyzed is indicated below each graph from 7 cotyledons, each from a different seedling. Similar results were obtained in 4 independent experiments. Scale bar = 50 µm, *t*-test **p<0.01.

(D) Absence of PC interdigitation gradient in the *tir1Qt*. (Left) Confocal image showing PI-stained epidermis of the entire cotyledon for 3-days-old wild type and *tir1Qt* seedlings. (Right) Heatmap of Margin Roughness (MR) values identified for each cell. High-low MR values are indicated by the orange-to-blue color gradient on the heatmap. Scale bar = 100 µm.

(E) PC shapes were analyzed for several gain-of-function mutants in the AUX/IAA family. The gain-of-function mutations for *AUX/IAA* genes are point mutations in the domain II of AUX/IAA proteins, causing their stabilization and inhibition of auxin transcriptional responses. Quantitative analysis of mean lobe numbers in cotyledon PCs from the *axr2-1*, *axr3- 1*, *shy2-3,* and *iaa18*$^{D}$ gain-of-AUX/IAA-function mutants. Both *iaa18*$^{D}$ and *shy2-3* showed a significant reduction in mean lobe number per cell area (*** p<0.001, *t*-test), with *iaa18*$^{D}$ having a stronger phenotype and was thus chosen for further study.

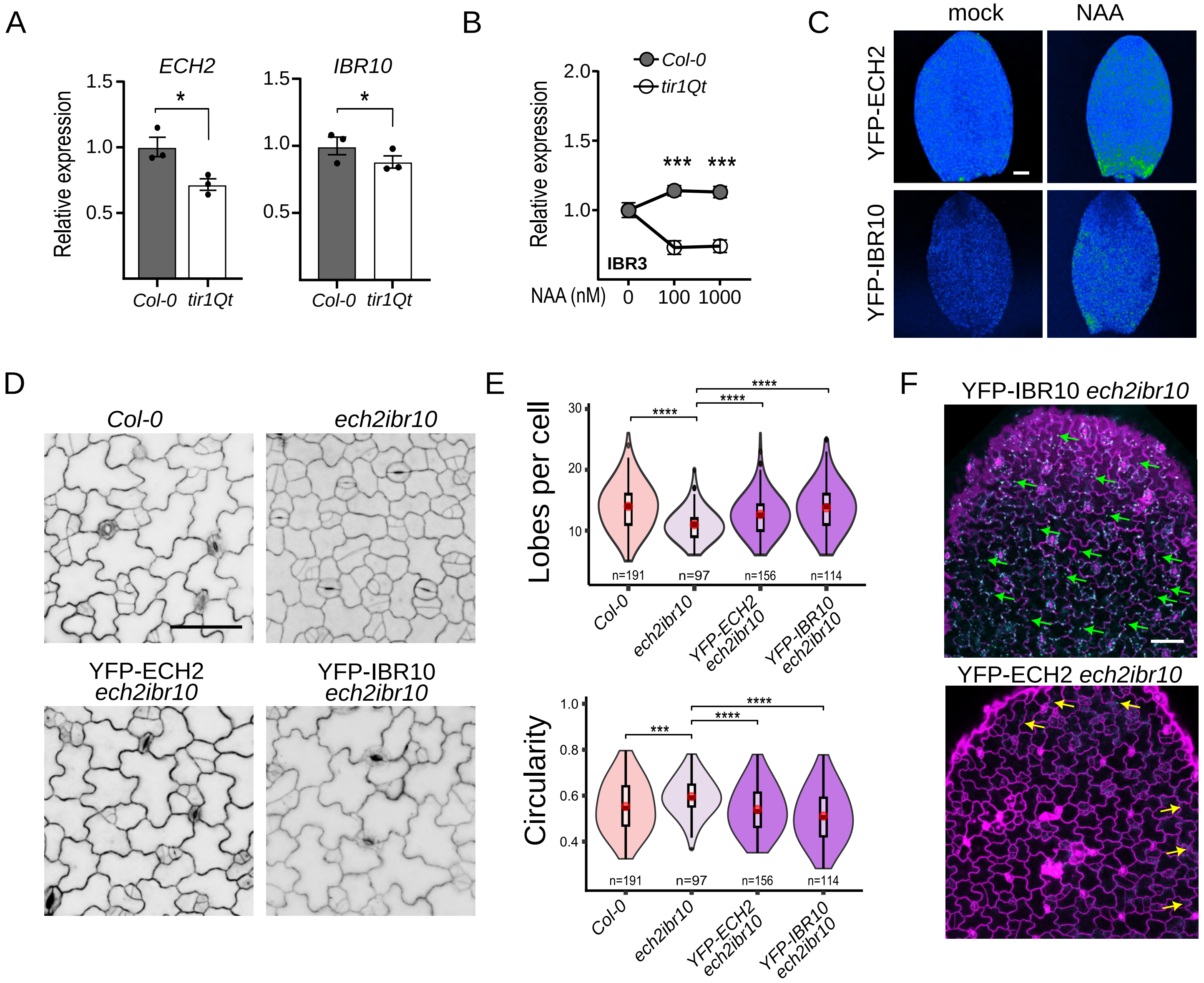
A
ECH2
IBR10
Relative expression
1.5
1.0
0.5
*
Col-0
tir1Qt
B
Col-0
tir1Qt
2.0
1.5
1.0
Relative expression
***
***
IBR3
NAA (nM)
0
100
1000
C
mock
NAA
YFP-ECH2
YFP-IBR10
D
Col-0
ech2ibr10
YFP-ECH2 ech2ibr10
YFP-IBR10 ech2ibr10
E
Lobes per cell
30
20
10
****
n=191
n=97
n=156
n=114
Col-0
ech2ibr10
YFP-ECH2 ech2ibr10
YFP-IBR10 ech2ibr10
Circularity
1.0
0.8
0.6
0.4
***
F
YFP-IBR10 ech2ibr10
YFP-ECH2 ech2ibr10

**Figure S5. The TIR1/AFB pathway promotes the expression of *ECH2/IBR* auxin biosynthetic genes that function in PC morphogenesis.**

(A) *ECH2* and *IBR10* expression was reduced in *tir1Qt*. qRT-PCR results of expression levels are relative to values obtained in wild-type Col-0 and normalized to the calibrator *UBQ5* (AT3G62250). RNA samples were obtained from ~30 *tir1Qt* class III dissected cotyledons from 4.5-old seedlings. Each dot in the graph represents a biological replicate, each with 3 technical replicates. *t*-test, *p<0.05.

(B) Auxin-induced IBR3 expression was compromised in *tir1Qt*. qRT-PCR results for *IBR3* expression levels were normalized to the geometric mean of the calibrators *PP2AA3* (AT1G13320), *AP4M* (AT4G24550), and relative to values obtained in mock conditions for each genotype. Samples from a pool of ~30 wild type (Col-0) and *tir1Qt* class III 4.5-days-old seedlings are treated for 0.5h in a liquid medium with the indicated concentration of auxin NAA. The graph shows the average of 3 biological replicates each with 3 technical replicates + SEM. *t*-test, ***p<0.001.

(C) Increased ECH2 and IBR10 protein accumulation in response to auxin. *ECH2::YFP-ECH2* and *IBR10::YFP-IBR10* seedlings (36 HAP without seedcoats) were treated with 10 µM NAA for 20 h in liquid medium, cotyledons dissected and imaged in z-stack. Maximum projections are shown. Similar results were obtained with 100 nM and 1 µM NAA. 6 cotyledons were analyzed per condition. These images are representative of 3 independent replicates. Scale bar = 50 µm.

(D) *YFP-ECH2 and YFP-IBR10* complement the PC phenotype of the double mutant *ech2-/-;ibr10-/-*. *ECH2::YFP-ECH2;ech2ibr10* and *IBR10::YFP-IBR10;ech2i*br10 seedlings were grown for 3 days in solid media, stained with 50 µg/mL propidium iodide for 30 seconds and imaged in the cotyledon's center with confocal microscopy. These images are representative of 3 independent experimental replicates. Scale bar = 50 µm.

(E) Quantification of data shown in D. Images were processed with PaCeQuant for automated segmentation and PC feature analysis. Lobes per cell and circularity (also called roundness or isoperimetric quotient). The sample size, n=97-156 cells, is indicated below each violin plot, coming from 6-11 different cotyledons, each from different seedlings. *t*-test, ***p<0.001, **** p<0.0001

(F) Representative images of *ECH2::YFP-ECH2*/*ech2ibr10* and *IBR10::YFP-IBR10*/*ech2ibr10* cotyledons displaying cell borders stained with propidium iodide (purple) and the respective fusion protein (cyan). Green arrows indicate YFP-IBR10 throughout the cotyledon's epidermis. Yellow arrowheads indicate YFP-ECH2 at the cotyledon's borders and tip. Scale bar = 50 µm.

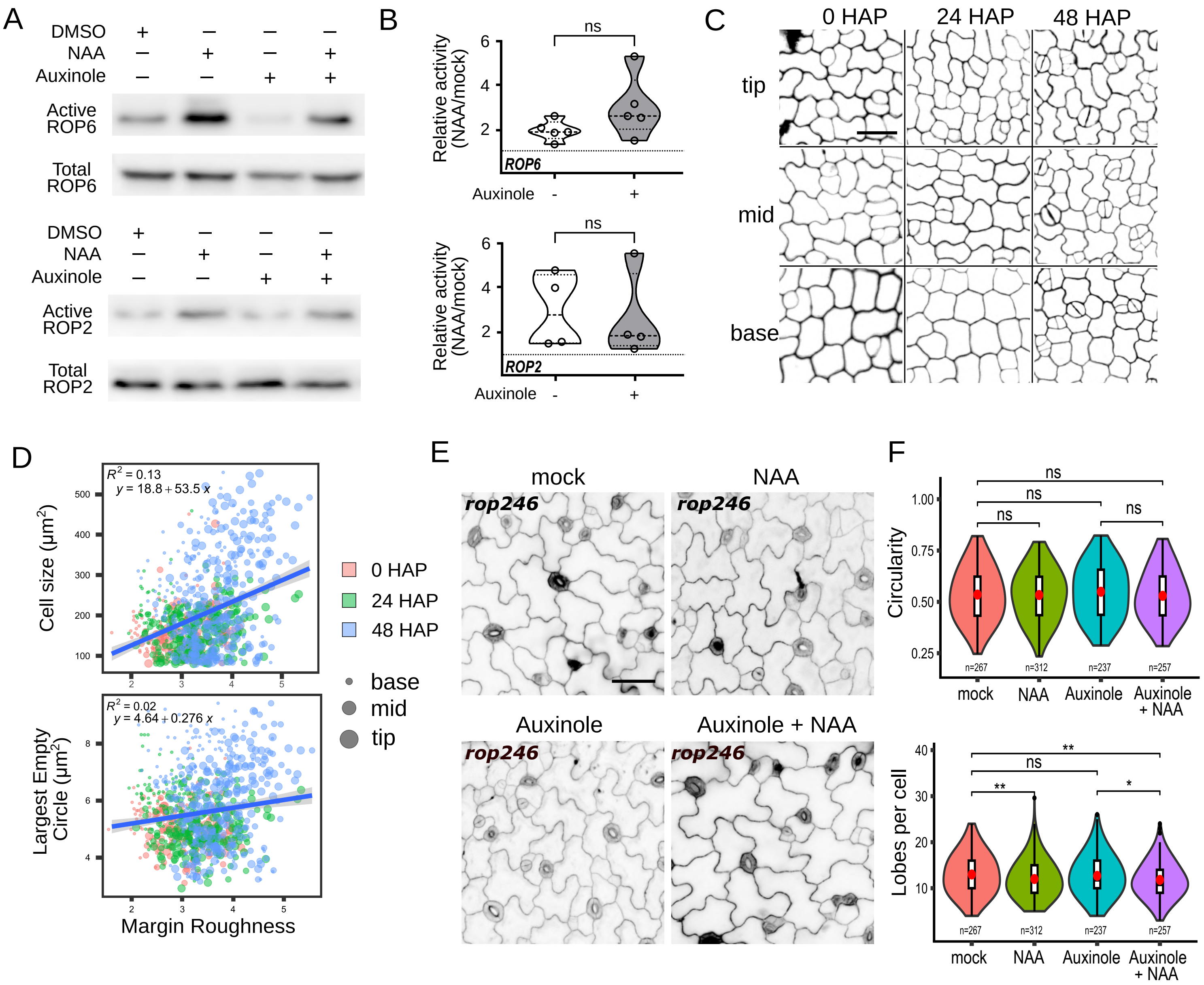
A
DMSO
NAA
Auxinole
Active ROP6
Total ROP6
Active ROP2
Total ROP2
B
Relative activity (NAA/mock)
ns
ROP6
ROP2
Auxinole
C
0 HAP
24 HAP
48 HAP
tip
mid
base
D
R2 = 0.13
y = 18.8 + 53.5 x
Cell size (µm2)
R2 = 0.02
y = 4.64 + 0.276 x
Largest Empty Circle (µm2)
Margin Roughness
0 HAP
24 HAP
48 HAP
base
mid
tip
E
mock
NAA
Auxinole
Auxinole + NAA
rop246
F
Circularity
ns
Lobes per cell
**
*
n=267
n=312
n=237
n=257
mock
NAA
Auxinole
Auxinole + NAA

**Figure S6. The promotion of PC interdigitation by TIR1/AFB-generated auxin is decoupled from cell expansion-induced mechanical stress.**

(A) Auxinole did not affect the induction of ROP2/ROP6 activity by auxin. GFP-ROP2 and GFP-ROP6 seedlings were grown in liquid media for 4 days and for 2 additional days in the presence of 200 μM auxinole in order to completely inhibit TIR1-AFB signaling. Then, cultured seedlings (about 20 seedlings per well) were treated with 1 μM NAA for 5 minutes before proteins were extracted for ROP activity assay by western blotting. The levels of total GFP-ROP2 and GFP-ROP6 and active (GTP-bound) GFP-ROP2 and GFP-ROP6 were analyzed using anti-GFP antibody.

(B) Quantification of auxin-induced ROP2/6 activity relative to mock conditions in wild-type Col-0 seedlings grown in the absence (white) or presence (gray) of auxinole shows that auxinole treatment did not affect the induction of ROP activity by auxin, indicating that the TIR1/AFB pathway acts upstream of the induction of TMK-dependent ROP activation. ROP activity is the ratio between active to total ROP. Relative ROP activity is the auxin treatments compared to mock conditions. Numbers above the dashed line at relative activity = 1, indicate auxin-induced ROP activity. Each dot represents a biological replicate, *t*-test, ns=not significant ($p$-values>0.05).

(C) Representative PC images from cells at the tip, middle, and base of cotyledons at 0, 24, and 48 HAP. Scale bar = 20 μm. Representative from 8 cotyledons, each from a different seedling.

(D) Interdigitation correlates weakly with cell size and does not correlate with mechanical stress. Multivariable correlation plots are shown for Margin Roughness (a proxy for interdigitation), cell size, and largest empty circle (a proxy for mechanical stress) at the tip, middle, and base of cotyledons 0, 24, and 48 HAP, obtained from images as shown in C. This is the aggregated data shown in Figure 6A to show that the weak correlation between interdigitation and cell size is due mainly to cells localized at the tip of 48 HAP cotyledons (big blue dots above the linear model line). The blue line represents the linear model. The gray shadow associated with the linear model represents the interval of 95% confidence. The equation for the linear model is on the upper left. The correlation coefficient ($R^2$) is shown above the linear equation.

(E) Representative PC images from 3 days-old *rop246* cotyledons treated for 2 days with 100 nM NAA in the absence or presence of 20 μM auxinole. Scale bar = 50 μm.

(F) Quantification of the *rop2 rop4 rop6* triple mutant (*rop246)* in auxin treatments shown in E. Seedlings were grown for 3 days in a liquid medium and then with 100 nM NAA for 48 h in the presence or absence of 200 μM auxinole. *Top panel*. Circularity (a proxy for interdigitation) is completely unaffected by treatments and co-treatments. *Bottom panel* - Lobes per cell are not increased by auxin in *rop246*, as reported previously [28]. Even more, *rop246* shows a slight decrease in lobe number suggesting there was isotropic growth without lobing. n = 237-312 cells from 15-19 cotyledons, each from a different seedling. *t*-test, *p<0.05, **p<0.01, ns =no significance.

### References


1. Ulmasov, T., Murfett, J., Hagen, G., and Guilfoyle, T.J. (1997). Aux/IAA proteins repress expression of reporter genes containing natural and highly active synthetic auxin response elements. Plant Cell *9*, 1963–1971. https://doi.org/10.1105/tpc.9.11.1963.

2. Lee, D.J., Kim, S., Ha, Y.-M., and Kim, J. (2008). Phosphorylation of Arabidopsis response regulator 7 (ARR7) at the putative phospho-accepting site is required for ARR7 to act as a negative regulator of cytokinin signaling. Planta *227*, 577–587. https://doi.org/10.1007/s00425-007-0640-x.

3. Yu, Y., Tang, W., Lin, W., Li, W., Zhou, X., Li, Y., Chen, R., Zheng, R., Qin, G., Cao, W., et al. (2023). ABLs and TMKs are co-receptors for extracellular auxin. Cell *186*, 5457-5471.e17. https://doi.org/10.1016/j.cell.2023.10.017.

4. Li, Q., Yang, J., Zhang, Y., Wang, F., Chang, M., Xu, T., Wang, S., and He, J. (2021). Transmembrane Kinases are essential for plant development. BioRxiv. https://doi.org/10.1101/2021.03.11.434919.

5. Xu, T., Dai, N., Chen, J., Nagawa, S., Cao, M., Li, H., Zhou, Z., Chen, X., De Rycke, R., Rakusová, H., et al. (2014). Cell surface ABP1-TMK auxin-sensing complex activates ROP GTPase signaling. Science *343*, 1025–1028. https://doi.org/10.1126/science.1245125.

6. Prigge, M.J., Platre, M., Kadakia, N., Zhang, Y., Greenham, K., Szutu, W., Pandey, B.K., Bhosale, R.A., Bennett, M.J., Busch, W., et al. (2020). Genetic analysis of the Arabidopsis TIR1/AFB auxin receptors reveals both overlapping and specialized functions. eLife *9*. https://doi.org/10.7554/eLife.54740.

7. Schepetilnikov, M., Makarian, J., Srour, O., Geldreich, A., Yang, Z., Chicher, J., Hammann, P., and Ryabova, L.A. (2017). GTPase ROP2 binds and promotes activation of target of rapamycin, TOR, in response to auxin. EMBO J. *36*, 886–903. https://doi.org/10.15252/embj.201694816.

8. Li, H., Xu, T., Lin, D., Wen, M., Xie, M., Duclercq, J., Bielach, A., Kim, J., Reddy, G.V., Zuo, J., et al. (2013). Cytokinin signaling regulates pavement cell morphogenesis in Arabidopsis. Cell Res. *23*, 290–299. https://doi.org/10.1038/cr.2012.146.

9. Brunoud, G., Wells, D.M., Oliva, M., Larrieu, A., Mirabet, V., Burrow, A.H., Beeckman, T., Kepinski, S., Traas, J., Bennett, M.J., et al. (2012). A novel sensor to map auxin response and distribution at high spatio-temporal resolution. Nature *482*, 103–106. https://doi.org/10.1038/nature10791.

10. Liao, C.-Y., Smet, W., Brunoud, G., Yoshida, S., Vernoux, T., and Weijers, D. (2015). Reporters for sensitive and quantitative measurement of auxin response. Nat. Methods *12*, 207–210, 2 p following 210. https://doi.org/10.1038/nmeth.3279.

11. Tian, Q., Uhlir, N.J., and Reed, J.W. (2002). Arabidopsis SHY2/IAA3 inhibits auxin-regulated gene expression. Plant Cell *14*, 301–319. https://doi.org/10.1105/tpc.010283.

12. Ploense, S.E., Wu, M.-F., Nagpal, P., and Reed, J.W. (2009). A gain-of-function mutation in IAA18 alters Arabidopsis embryonic apical patterning. Development *136*, 1509–1517. https://doi.org/10.1242/dev.025932.

13. Damodaran, S., and Strader, L. (2023). Mechanisms governing competence for cell reprogramming in arabidopsis adventitious root formation. https://doi.org/10.2139/ssrn.4392154.

14. Kusaka, N., Maisch, J., Nick, P., Hayashi, K., and Nozaki, H. (2009). Manipulation of intracellular auxin in a single cell by light with esterase-resistant caged auxins. Chembiochem *10*, 2195–2202. https://doi.org/10.1002/cbic.200900289.

15. Hayashi, K., Neve, J., Hirose, M., Kuboki, A., Shimada, Y., Kepinski, S., and Nozaki, H. (2012). Rational design of an auxin antagonist of the SCF(TIR1) auxin receptor complex. ACS Chem. Biol. *7*, 590–598. https://doi.org/10.1021/cb200404c.

16. Schindelin, J., Arganda-Carreras, I., Frise, E., Kaynig, V., Longair, M., Pietzsch, T., Preibisch, S., Rueden, C., Saalfeld, S., Schmid, B., et al. (2012). Fiji: an open-source platform for biological-image analysis. Nat. Methods *9*, 676–682. https://doi.org/10.1038/nmeth.2019.

17. Möller, B., Poeschl, Y., Plötner, R., and Bürstenbinder, K. (2017). PaCeQuant: A Tool for High-Throughput Quantification of Pavement Cell Shape Characteristics. Plant Physiol. *175*, 998–1017. https://doi.org/10.1104/pp.17.00961.

18. Möller, B., Poeschl, Y., Klemm, S., and Bürstenbinder, K. (2019). Morphological Analysis of Leaf Epidermis Pavement Cells with PaCeQuant. Methods Mol. Biol. *1992*, 329–349. https://doi.org/10.1007/978-1-4939-9469-4_22.

19. Poeschl, Y., Möller, B., Müller, L., and Bürstenbinder, K. (2020). User-friendly assessment of pavement cell shape features with PaCeQuant: Novel functions and tools. Methods Cell Biol. *160*, 349–363. https://doi.org/10.1016/bs.mcb.2020.04.010.

20. Jefferson, R.A., Kavanagh, T.A., and Bevan, M.W. (1987). GUS fusions: beta-glucuronidase as a sensitive and versatile gene fusion marker in higher plants. EMBO J. *6*, 3901–3907. https://doi.org/10.1002/j.1460-2075.1987.tb02730.x.

21. Jefferson, R.A. (1989). The GUS reporter gene system. Nature *342*, 837–838. https://doi.org/10.1038/342837a0.

22. Blázquez, M. (2007). Quantitative GUS activity assay in intact plant tissue. CSH Protoc. *2007*, pdb.prot4688. https://doi.org/10.1101/pdb.prot4688.

23. Halder, V., and Kombrink, E. (2015). Facile high-throughput forward chemical genetic screening by in situ monitoring of glucuronidase-based reporter gene expression in Arabidopsis thaliana. Front. Plant Sci. *6*, 13. 10.3389/fpls.2015.00013.

24. Ramakers, C., Ruijter, J.M., Deprez, R.H.L., and Moorman, A.F.M. (2003). Assumption-free analysis of quantitative real-time polymerase chain reaction (PCR) data. Neurosci. Lett. *339*, 62–66. https://doi.org/10.1016/s0304-3940(02)01423-4.

25. Vandesompele, J., De Preter, K., Pattyn, F., Poppe, B., Van Roy, N., De Paepe, A., and Speleman, F. (2002). Accurate normalization of real-time quantitative RT-PCR data by geometric averaging of multiple internal control genes. Genome Biol. *3*, RESEARCH0034. https://doi.org/10.1186/gb-2002-3-7-research0034.

26. Hellemans, J., Mortier, G., De Paepe, A., Speleman, F., and Vandesompele, J. (2007). qBase relative quantification framework and software for management and automated analysis of real-time quantitative PCR data. Genome Biol. *8*, R19. https://doi.org/10.1186/gb-2007-8-2-r19.

27. Mustroph, A., Zanetti, M.E., Jang, C.J.H., Holtan, H.E., Repetti, P.P., Galbraith, D.W., Girke, T., and Bailey-Serres, J. (2009). Profiling translatomes of discrete cell populations resolves altered cellular priorities during hypoxia in Arabidopsis. Proc Natl Acad Sci USA *106*, 18843–18848. https://doi.org/10.1073/pnas.0906131106.

28. Xu, T., Wen, M., Nagawa, S., Fu, Y., Chen, J.-G., Wu, M.-J., Perrot-Rechenmann, C., Friml, J., Jones, A.M., and Yang, Z. (2010). Cell surface- and rho GTPase-based auxin signaling controls cellular interdigitation in Arabidopsis. Cell *143*, 99–110. https://doi.org/10.1016/j.cell.2010.09.003.
